# Tumor suppressor PLK2 may serve as a biomarker in triple-negative breast cancer for improved response to PLK1 therapeutics

**DOI:** 10.1101/2021.06.16.448722

**Authors:** Yang Gao, Elena B. Kabotyanski, Elizabeth Villegas, Jonathan H. Shepherd, Deanna Acosta, Clark Hamor, Tingting Sun, Celina Montmeyor-Garcia, Xiaping He, Lacey E. Dobrolecki, Thomas F. Westbrook, Michael T. Lewis, Susan G. Hilsenbeck, Xiang H.-F. Zhang, Charles M. Perou, Jeffrey M. Rosen

## Abstract

Polo-like kinase (PLK) family members play important roles in cell cycle regulation. The founding member PLK1 is oncogenic and preclinically validated as a cancer therapeutic target. Paradoxically, PLK2 (chromosome 5q11.2) is frequently deleted in human breast cancers, preferentially in basal-like and triple-negative breast cancer subtypes. Here, we found that PLK2 was tumor suppressive in breast cancer and knockdown of PLK1 rescued phenotypes induced by PLK2-loss both *in vitro* and *in vivo*. We also demonstrated that PLK2 directly interacted with PLK1 at prometaphase and that mutations in the kinase domain of PLK2, but not polo-box binding domains, changed their interaction pattern. Furthermore, treatment of syngeneic transplantation mouse tumor models and patient-derived xenografts using the PLK1 inhibitor volasertib alone, or in combination with carboplatin, indicated that PLK2-low breast tumors had a significantly better response to these drugs. Re-expression of PLK2 in an inducible PLK2-null mouse model reduced the therapeutic efficacy of volasertib. Taken together, our data suggest PLK2 loss may serve as a biomarker to predict response to PLK1 therapeutics, alone and in combination with chemotherapy.

**Significance:** The tumor suppressive role of PLK2, and its relationship with the oncogene PLK1, provide a mechanistic rationalization to use PLK1 inhibitors in combination with chemotherapy to treat PLK2 low/deleted tumors. TNBC, and other cancers with low PLK2 expression, are such candidates to leverage precision medicine to identify patients who might benefit from treatment with these inhibitors.

## Introduction

Breast cancer remains the most prevalent cancer and the second leading cause of cancer deaths in American women in 2021 (1). Therefore, it is imperative to understand the molecular mechanisms underlying breast cancer development in order to develop novel therapies. Categorization of breast cancer based on its status of estrogen receptor (ER), progesterone receptor (PR), and human epidermal growth factor receptor 2 (HER2) has markedly improved the treatment of the luminal and HER2-specific subtypes (2, 3). For example, the luminal subtype (ER+, PR±) typically responds to endocrine therapies, and the HER2 clinical subtype (HER2+, ER-/+, PR-/+) is treated with the HER2-targeted drug trastuzumab (4–8). However, targeted therapies are still in urgent need for triple-negative breast cancer (TNBC, ER-, PR-, and HER2-) due to the lack of these standard therapeutic targets. In this study, we identified a potent candidate biomarker, Polo-like kinase 2 (PLK2, chromosome 5q11.2), and a possible therapeutic target (PLK1), that may not only help stratify TNBC patients for treatment, but also be relevant for patients with several other cancers that exhibit chromosome 5q loss across the PLK2 region (9, 10).

PLK2 belongs to a gene family consisting of five serine/threonine kinases (PLK1-PLK5) with highly conserved N-terminal kinase domains and C-terminal polo-box domains (PBDs). PLK family members play important roles in regulating the cell cycle and DNA damage response (11–13). The founding and most well-studied member of the family, PLK1, is expressed in the proliferating cells of normal tissues with a peak during G2/M and controls many cell cycle events including centrosome maturation, mitotic entry, chromosome segregation, and cytokinesis (13–15). Extensive studies have shown that PLK1 overexpression is oncogenic in many types of cancer and it has been validated preclinically as a cancer therapeutic target (16–18). Several PLK1 inhibitors, such as volasertib (BI 6727) (19), are currently under clinical development. Volasertib plus carboplatin has demonstrated activity in heavily pretreated patients with advanced solid tumors (20).

PLK2 was initially named serum-inducible kinase (SNK) because it is an early growth response gene upon serum treatment (21). PLK2 is mainly localized in centrosomes during the G1 phase and is necessary for centriole duplication (22–24). It also functions in the nervous system by regulating homeostatic synaptic plasticity and neuronal cell differentiation (13,25,26). Plk2 null mice display embryonic growth retardation but are viable, which may be due to compensatory effects of other Plks (27). PLK2 is highly expressed in the mammary gland (28). In contrast to *in vitro* studies, deletion of Plk2 in mouse mammary epithelial cells (MECs) *in vivo* surprisingly not only induces cell proliferation and gland hyperbranching, but also disrupts mitotic spindle orientation and cellular polarity (28, 29). Moreover, loss of Plk2 results in an increased number of less differentiated pre-neoplastic lesions in the mammary glands of multiparous mice, suggesting PLK2 is required for normal mammary gland development (29).

Although PLK2 belongs to the same family as PLK1, and both kinases are implicated in cell cycle progression, recent studies as well as our data presented herein, suggest that PLK2 may actually be a tumor suppressor that is silenced in many types of cancers, including breast cancer (30–34). This paradox stimulated us to investigate the functions of PLK1 and PLK2 in breast cancer, as well as potential clinical implications. Here, we demonstrated that PLK2 functions as a tumor suppressor in breast cancer, and suggested that its tumor suppressive role is mediated, at least partially, by its direct interaction with PLK1. Furthermore, using both preclinical genetically engineered mouse (GEM) models and patient-derived xenograft (PDX) models, we showed that PLK2 loss may serve as a potential biomarker to predict response to PLK1 inhibitors, alone and in combination with chemotherapy.

## Results

### PLK2 is a tumor suppressor in breast cancer

Previous studies from our laboratory and others have suggested that there is a frequent arm-level loss of chromosome 5q region across different types of cancer, including breast cancer, lung squamous cell carcinoma, and ovarian cancer [(9, 10) and Siegel, Perou, et al., manuscript in preparation]. To explore the gene-specific copy number alterations (CNAs) of PLK2 (chromosome 5q11.2), we analyzed the TCGA Pan-Cancer and Tumorscape data sets using the TCGA Copy Number Portal from the Broad Institute and confirmed that PLK2 is significantly deleted in several types of cancer, including breast cancer (Figure 1A and Supplemental Figure S1A). This result is consistent with our previous report that basal-like breast tumors have a frequent loss of chromosome 5q11-35 that typically included PLK2 and the checkpoint clamp loader component gene RAD17 (35). To gain more insight into the CNAs in different breast cancer subtypes, we performed GISTIC analysis on TCGA breast tumors and discovered that the loss of PLK2 was identified in 21.8% of all breast cancer patients (Figure 1B). Among these, loss of PLK2 was found in 11.3% of ER+, 24.4% of HER2+, but markedly in 58% of TNBC patients. Similar to PLK2, loss of RAD17 was also observed at a higher level in TNBC (Figure 1B). In contrast, the gain of PLK1, the founding member of the PLK family and a well-known oncogene, was less in the TNBC subtype (26.7% of TNBC vs. 42.9% of all, 46.9% of ER+, and 45.1% of HER2+, Figure 1B).

**Figure 1.**
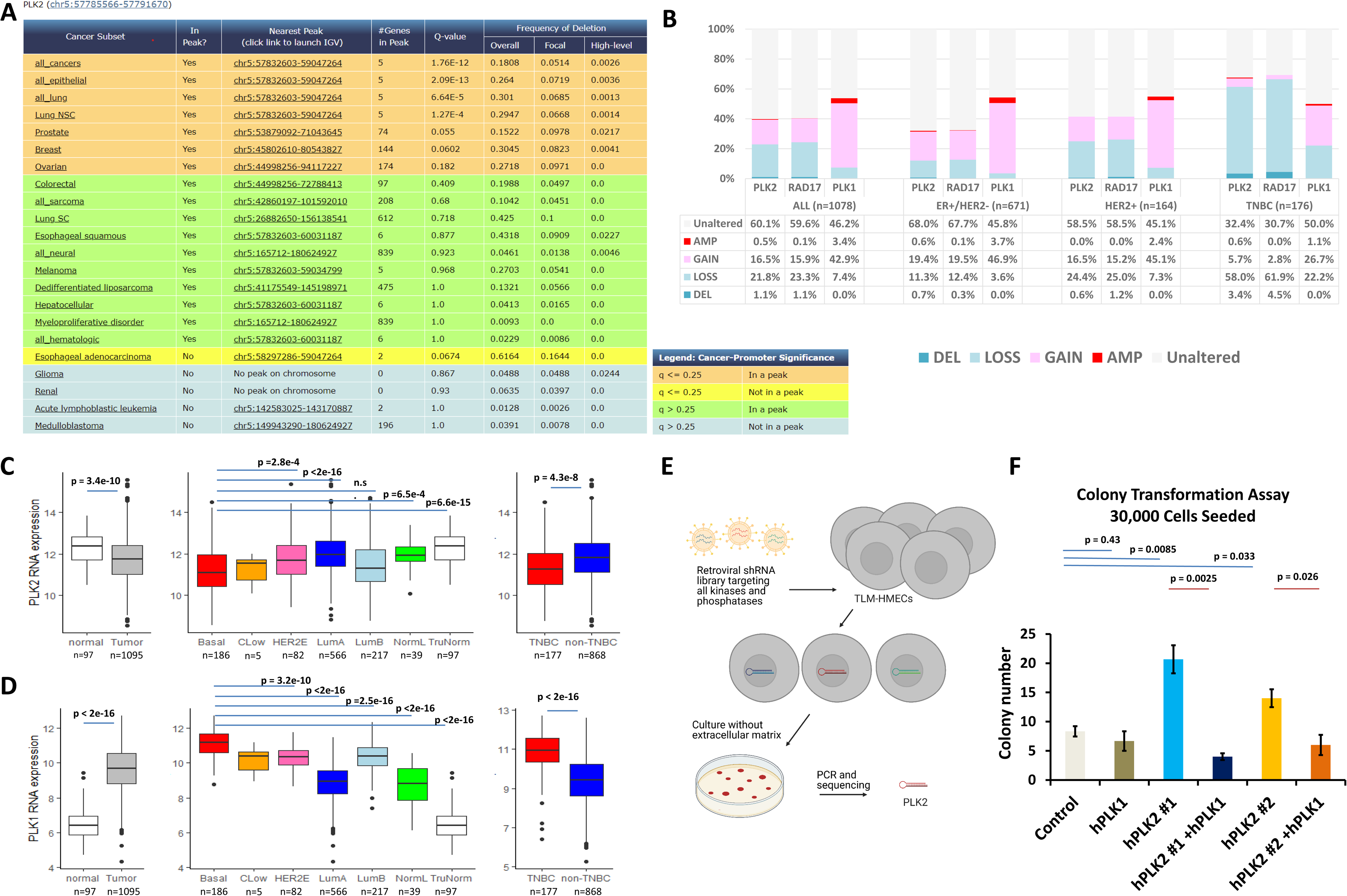
PLK2 is tumor suppressive and knockdown of PLK1 rescues PLK2 loss induced transformation of HMECs. **(A)** PLK2 is significantly deleted in 5 of 16 independent cancer types (q<=0.25) in the Tumorscape dataset (2011-02-01). Among these, PLK2 is located within a peak region of deletion in 4 cancer types including non-small cell lung cancer, prostate cancer, breast cancer, and ovarian cancer. **(B)** PLK2, RAD17, and PLK1 copy number alterations in different breast cancer subtypes. Note that the loss of PLK2 and RAD17 is more dramatic in the TNBC subtype as compared to others, but the gain of PLK1 is less prominent in the TNBC subtype. Amplification (Amp) >2.0, Gain >1.0, Deletion (Del) <2.0, Loss <1.0. PLK2 **(C)** and PLK1 **(D)** RNA expression vary significantly between normal breast tissue and breast tumor samples (left), intrinsic breast cancer subtypes (center), and triple-negative (TNBC) versus non-TNBC patients from the TCGA breast tumor dataset. P-values shown from student’s t-test for two-class comparisons (normal vs Tumor and TNBC vs non-TNBC); P-values between intrinsic subtypes are pairwise t-tests with Bonferroni correction for multiple comparisons. **(E)** Schematic of the unbiased RNAi-based forward genetic screen. A library of retroviral shRNAs targeting all human kinases and phosphatases was transduced into TLM-HMECs in duplicate. Five hundred and thirty anchorage-independent macroscopic colonies were quantitated from two independent screens. Colonies containing shRNAs were identified by PCR amplification and sequencing. **(F)** Knockdown of PLK2 using two shRNAs increased the colony number for both, suggesting a tumor suppressive role of PLK2 in HMECs. This induction was abolished by additional knockdown of PLK1. PLK1 shRNA alone didn’t show an effect on colony number formed in the colony transformation assay. Thirty thousand HMECs were seeded in the plates. Assays were performed in triplicate. Statistical significance was determined using unpaired student’s t-test.

We then examined mRNA expression in the TCGA database to further investigate the relationship between PLK2 and PLK1 in breast cancer. Patients with primary breast tumors had lower expression of PLK2 mRNA, but higher levels of PLK1 mRNA, relative to normal breast tissue (Figure 1C and 1D). Further analysis indicated that lower expression of PLK2 and a higher level of PLK1 mRNA was specifically linked to the basal-like subtype or TNBC subtype (Figure 1C and 1D, Supplemental Figure S1B and S1C).

In support of the functional consequences of these human tumor findings, an unbiased RNAi-based forward genetic screen also identified PLK2 as one of the candidate tumor suppressors in breast cancer (Figure 1E) (36, 37). In brief, human mammary epithelial cells (HMECs) were immortalized by transducing with human telomerase catalytic subunit (hTERT) and SV40 large T-antigen (LT), hereby designated TLM-HMECs. These cells, however, need to be anchored to the extracellular matrix (ECM) to proliferate. Therefore, by applying an shRNA library that targeted all kinases and phosphatases to TLM-HMECs cultured in the absence of ECM, we were able to discover critical candidate genes involved in cell transformation (37), including PLK2. Subsequently, we performed a colony transformation assay and found that knockdown of PLK2 in TLM-HMECs using two different shRNAs, both increased the number of colonies as compared to controls (Figure 1F and Supplemental Figure S1D to S1F), consistent with the tumor suppressive role of PLK2.

### PLK1 mediates PLK2-loss induced phenotypes

PLK1 is overexpressed in breast cancer, especially TNBC, and targeting PLK1 has been reported to impair TNBC growth (38, 39). The apparent paradox that PLK1 and PLK2 might exert opposite roles in breast cancer led us to study potential links between PLK2 and PLK1. First, we tested the effect of PLK1 knockdown using a human PLK1 shRNA in the TLM-HMEC colony transformation assay. Knockdown of PLK1 alone did not significantly change the number of colonies formed, but instead it mitigated the increased colony formation observed following PLK2 knockdown, thus suggesting that the tumor suppressive function of PLK2 is specifically related to PLK1 activity (Figure 1G).

Next, we sought to examine whether inhibition of PLK1 was able to rescue the phenotypes observed following PLK2 loss in the mouse mammary gland *in vivo*. We isolated MECs from 8-week-old Plk2^-/-^ mice and transduced them with lentiviral vectors containing control, mouse mPlk1 #1, or mPlk1 #2 shRNA (Supplemental Figure S2). The cells were then injected into immunocompromised mice with cleared mammary fat pads. Transplanted outgrowths were harvested and characterized 8 weeks post-transplantation (Figure 2A). Visualization of the fluorescent whole-mounts and carmine alum staining showed that, as expected, the control outgrowths had increased branching, a phenotype previously observed in the Plk2^-/-^ mammary glands (29). Interestingly, the hyperbranched phenotype observed in the control appeared to be absent in the Plk1 knockdown outgrowths. Quantification of the branchpoints per millimeter of duct revealed that the controls had about 2 branchpoints per millimeter of duct, whereas both the mPlk1 #1 and mPlk1 #2 shRNA knockdown groups had about 1.4 branchpoints per millimeter, suggesting Plk1 knockdown alleviated the hyperbranching phenotype in Plk2^-/-^ mammary gland (Figure 2B).

**Figure 2.**
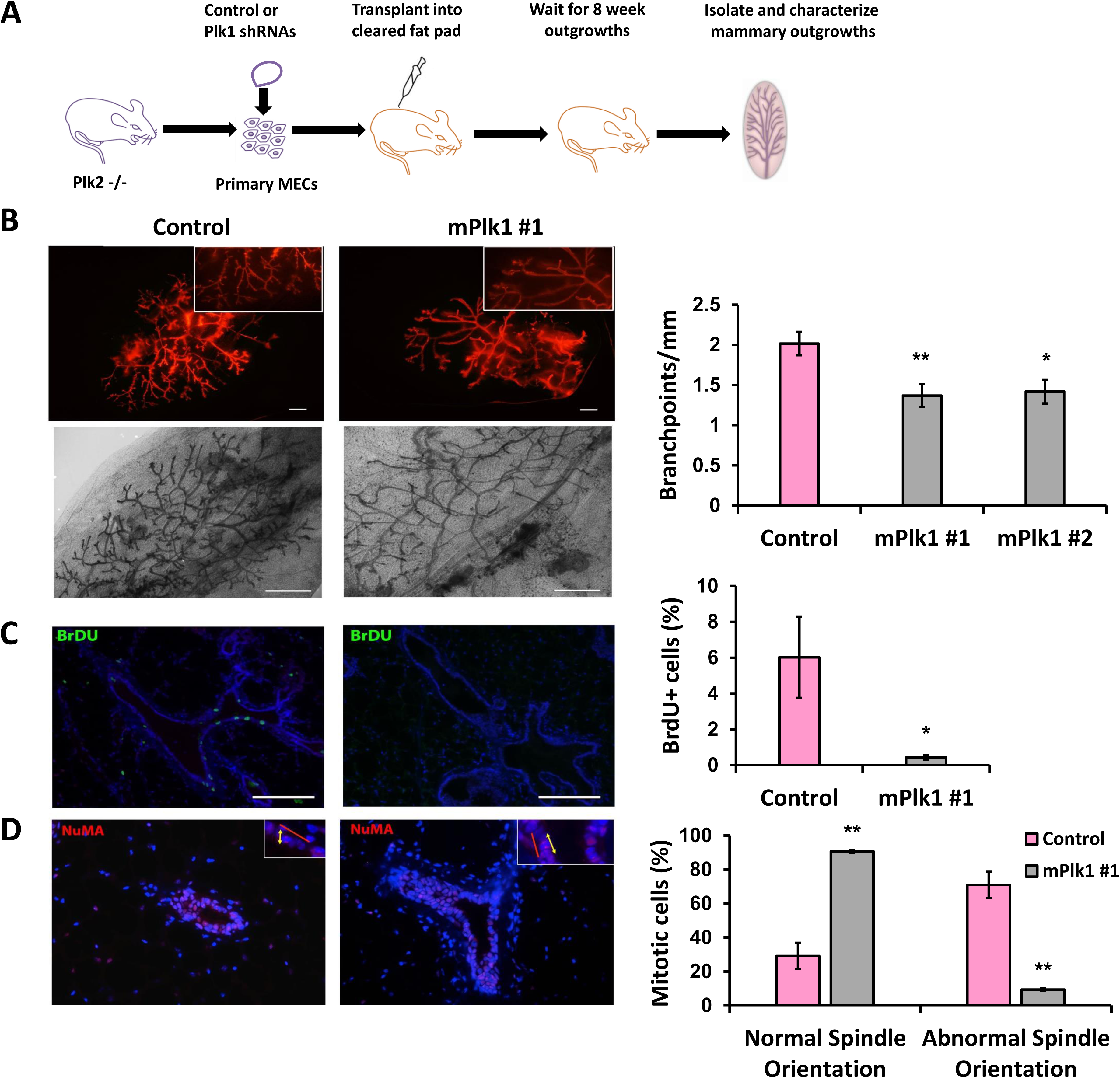
Knockdown of Plk1 rescues the Plk2^-/-^ mammary gland phenotypes. **(A)** Schematic representation of the experimental procedure used to determine the effect of Plk1 shRNA knockdown on Plk2^-/-^ MECs. **(B)** Whole-mount analyses of mammary glands transduced with control or Plk1 shRNAs showed a decrease in branching upon depletion of Plk1. n = 10 mice for Control group, n = 3 mice for mPlk1 #1 group, n = 7 mice for mPlk1 #2 group. Td-tomato red expression in the transduced cells is shown in the whole-mounts (Upper left panel). The lower left panel is stained with carmine alum. Representative pictures of mPlk1 #1 shRNA group were shown. The right panel shows the quantitation of branch points per millimeter. **(C)** Immunofluorescence for BrdU (green) incorporation and DAPI (blue) on paraffin-embedded sections of mammary glands demonstrated that knockdown of Plk1 abolished the hyperproliferation phenotype of Plk2^-/-^ MECs [30] (Left panel). n = 3 mice per group. The right panel shows the quantitation of BrdU positive MECs. **(D)** Disruption of mitotic spindle orientation in Plk2^-/-^ MECs was rescued by knockdown of Plk1. Normal Spindle orientation was defined as the angle between the basement membrane and the plane of the mitotic spindle being 0-10°, while abnormal being 10-90°. Mice were treated with estrogen and progesterone for two days to induce epithelial cell proliferation. n = 3 mice per group. Nuclear mitotic apparatus protein 1 (NuMA) (red) was used to stain the mitotic spindle. DAPI (blue) counterstained the nucleus. The red line indicates the proper plane of cell division and the yellow arrows denote the actual plane of division. The right panel shows the quantitation of normal and abnormal spindles. Statistical significance was determined by unpaired student’s t-test compared to the control group. *, p < 0.05; **, p < 0.01.

Mammary gland ductal epithelial cells have very low levels of proliferation when female mice become mature and the mammary ducts reach the edge of the mammary fat pad. However, Plk2 inactivation in mammary ducts leads to hyperproliferation and disorientated mitotic spindle formation in luminal epithelial cells, which exhibited an angle greater than 10° to the basement membrane (29). Subsequent knockdown of Plk1 decreased the high level of proliferating epithelial cells [∼6% bromodeoxyuridine (BrdU) positive] to the level usually observed in ducts in mature mice (∼0.4% BrdU positive) (Figure 2C). In addition, when the mice were treated with estrogen and progesterone to induce proliferation, knockdown of Plk1 decreased the percentage of mitotic epithelial cells with misoriented mitotic spindles and increased the presence of mitotic cells with normal spindle orientation parallel to the basement membrane (Figure 2D). These data suggest that the phenotypes associated with PLK2 loss are mediated, at least partially, by PLK1 during mammary gland development.

### PLK2 directly interacts with PLK1

Since homozygous deletion of Plk1 (Plk1^-/-^) results in embryonic lethality, but Plk1 heterozygotes (Plk1^+/-^) and Plk2 null (Plk2^-/-^) mice are viable (27, 40), we then attempted to breed Plk2^-/-^ with Plk1^+/-^ to determine if we could genetically rescue the Plk2^-/-^ phenotypes in mammary epithelial cells. Despite numerous matings, we were, however, unable to generate any offspring with the Plk2^-/-^; Plk1^+/-^ genotype (Supplemental Table S1). Further studies are required to determine the precise cause, but the synthetic lethality during embryonic development induced by the simultaneous loss of Plk2 and Plk1 suggested that there might be a genetic interaction between these two PLK family members.

Next, to examine whether PLK2 directly binds with PLK1, we performed a bimolecular fluorescence complementation (BiFC) assay. A wild-type PLK2 (bait) was fused with the N-terminus of yellow fluorescent protein (YFP) and the PLK1 (prey) was fused with the C-terminus of YFP. If an interaction occurs between PLK2 and PLK1, the two YFP fragments will come together and form a fluorescent YFP protein that can be detected by a flow cytometer (Figure 3A). Two previously identified substrates of PLK2, checkpoint kinase 1 (CHK1) and beta-tubulin (TUBB) were used as positive preys (41, 42). RB transcriptional corepressor 1 (RB1) was included as a negative control prey, which was shown previously not to interact with Plk2 using an immunoprecipitation pull-down assay (41). Bait and prey constructs were stably transduced into cells and analyzed for YFP fluorescence by flow cytometry 48 hours post-transduction. As expected, strong interactions between PLK2 and the positive controls, CHK1 and TUBB, were detected, while the negative control, RB1, showed a very low percentage of YFP-positivity (Figure 3B). More importantly, a significant interaction between PLK2 and PLK1 was observed (Figure 3B), thus indicating close physical proximity to one another in the cell.

**Figure 3.**
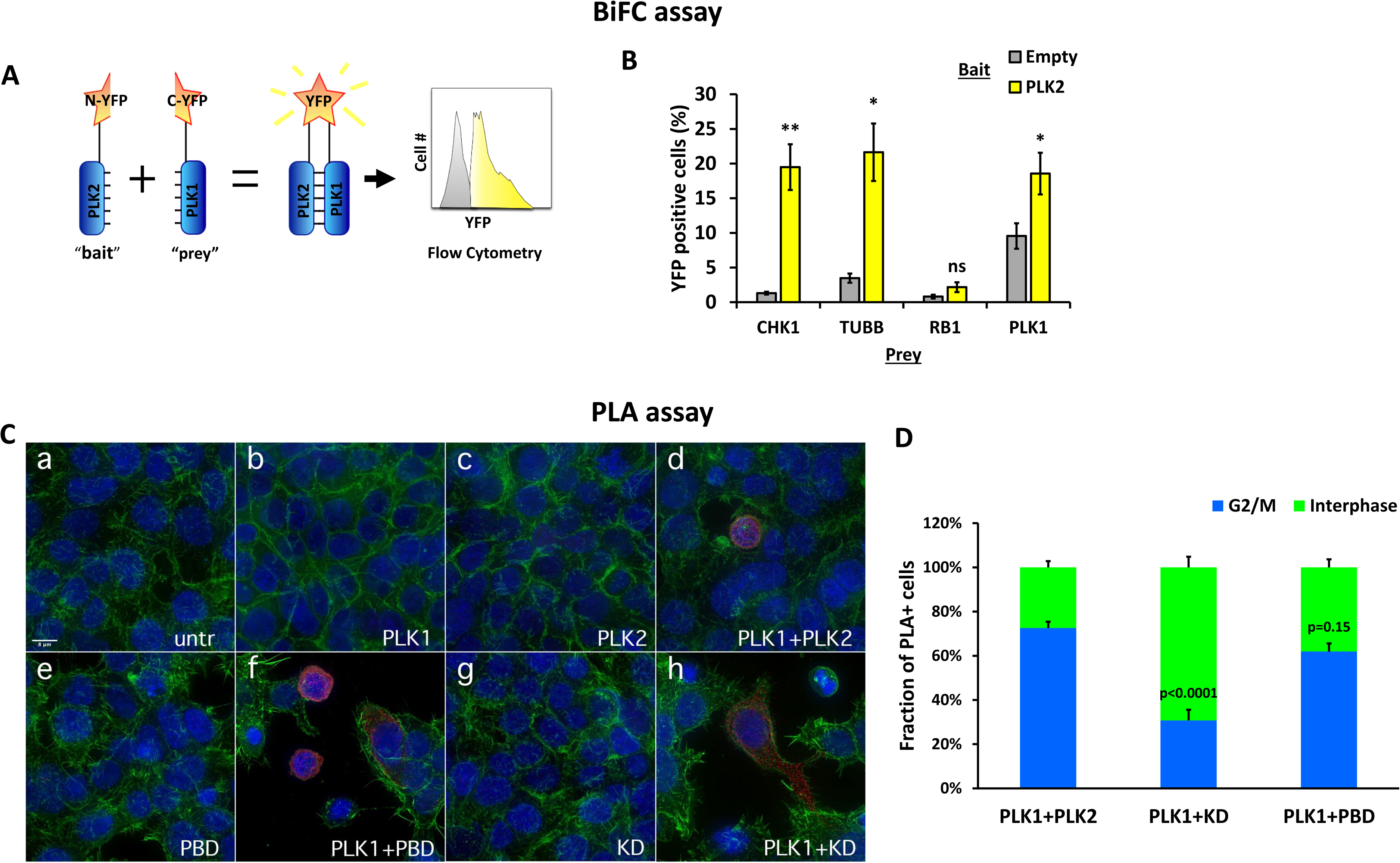
PLK2 directly interacts with PLK1. **(A)** Diagram of BiFC assay. The prey fused with the C-terminus of YFP is added to the bait fused with the N-terminus of YFP. If an interaction occurs between the prey and the bait, the two YFP fragments will come together and form a fluorescent YFP that can be analyzed by flow cytometry. **(B)** PLK2 directly interacted with PLK1 as well as CHK1 and TUBB. As predicted, no interaction was observed between PLK2 and RB1. Assays were repeated five times in triplicate using HEK293T cells. Statistical significance was determined by unpaired student’s t-test compared to the control group. Ns, not significant; *, p < 0.05; **, p < 0.01. **(C)** Proximity ligation assay in HEK293T cells co-expressing PLK1 with PLK2-wild type, PLK2-PBD, or PLK2-KD mutants. Red: PLA signals. Blue: DAPI nuclei staining. Green: ß-actin staining. Representative images with positive PLA signals for the interaction of PLK1 with PLK2 (image d), PLK1 with PLK2-PBD (polo-box domain mutant) (image f), and PLK1 with PLK2-KD (kinase-dead mutant) (image h) are shown. Negative controls: non-transfected cells (image a); cells expressing only one protein: PLK1 (image b), PLK2 (image c), PLK2-PBD (image e), or PLK2-KD (image g). Note that the PLA signals of PLK1+PLK2 and PLK1+PBD are located at the nucleus and/or perinuclear area of round G2/M cells while the PLA signals of PLK1+KD are located in the cytoplasm of flat and elongated interphase cells. **(D)** Quantitative analysis of PLA signals in cells co-expressing PLK1 with wild-type PLK2, PLK2-PBD, and PLK2-KD mutants using microscopy images. PLA-positive cells were divided into two groups: cells that have rounded morphology with condensed nuclear DNA were considered mitotic (G2/M), while cells with flat and elongated morphology and non-condensed nuclear DNA were considered to be in interphase of the cell cycle. PLA-positive cells (200 to 300) were counted per group. Statistical was determined by one-way ANOVA followed by Tukey test for multiple comparisons.

To confirm these findings at the single-cell level, we performed a complementary study using a proximity ligation assay (PLA) (Figure 3C and 3D). In HEK293T cells co-expressing both PLK1 and PLK2, we observed a strong positive PLA signal (red dots) as evidence for protein-protein interaction (Figure 3C, image d). As shown by confocal imaging, both proteins were at close proximity to each other in cells at the G2/M transition stage (presumably at the prometaphase stage), as evidenced by their rounded morphology and condensed nuclear DNA. Quantification revealed that 73% of 240 counted PLA-positive cells co-expressing PLK1 and PLK2 exhibited a rounded morphology, with the protein-protein interaction detected in the nucleus and/or perinuclear area (Figure 3D). The percentage of those cells was increased to 93% after treatment with nocodazole (40 ng/ml, 16 hr), which induced cells to arrest in prometaphase (Supplemental Figure S3B and S3C). No PLA signals were detected in non-transfected cells (Figure 3C, image a), or in cells expressing only one of the proteins (Figure 3C, images b and c). Additional negative controls were included for PLA performed in the absence of primary antibodies (Supplemental Figure S3A, images a to d). These results indicated that the PLK2-PLK1 interaction is cell cycle dependent.

It is well established that the polo-box domain and kinase domain of PLK1 play significant roles in proper subcellular localization and cell cycle, as well as PLK1-dependent protein-protein interactions (43). Therefore, we tested PLK2-PBD [polo-box domain mutant (W503F, H629A, K631M) at the C-terminus] and PLK2-KD [kinase-dead mutant (K111R) at the N-terminus] mutants in the context of PLK1-PLK2 interaction using PLA. Contrary to our expectations, mutations at the polo-box domain of PLK2 did not disrupt PLK1-PLK2 interactions, and cells co-expressing PLK1 and PLK2-PBD mutant revealed a similar pattern for positive PLA signals as compared to those observed for cells co-transfected with PLK1 and wild-type PLK2 (Figure 3C, image f). Most PLA signals were detected in the nucleus and perinuclear compartments, and 62% of 280 counted PLA-positive cells displayed rounded morphology with condensed DNA as a sign of early prometaphase (Figure 3D). Interestingly, in cells co-expressing PLK1 with PLK2-KD mutant, we observed positive PLA signals predominantly in the cytoplasm and 69% of 300 counted cells had a flat and elongated morphology with non-condensed DNA, clearly showing that interaction between the two proteins mostly occurs during the interphase but not the mitosis (Figure 3C, image h, and Figure 3D). In addition, the percentage of such cells did not significantly change following nocodazole treatment (Supplemental Figure S3B and S3C). These data suggest that the kinase domain of PLK2 most likely plays an important role in maintaining proper protein-protein interactions between PLK1 and PLK2 during the prometaphase stage of a cell cycle. No PLA signals were detected in cells expressing PLK2-PBD or PLK2-KD mutants alone (Figure 3C, images e and g), or when PLA was performed in the absence of primary antibodies (Supplemental Figure S3A, images e to h).

### Low PLK2 may serve as a biomarker to predict response to the PLK1 inhibitor

The previous combination of biochemical, genetic, and bioinformatic studies suggests that inhibiting PLK1 in a setting with low PLK2 expression, such as TNBC, may provide a targeted approach for a disease in which chemotherapy is still the only systematic treatment (44). Therefore, we tested the effects of a PLK1 inhibitor both as a single agent and in combination with chemotherapy in both novel p53^-/-^ GEM and PDX preclinical models with low vs. high PLK2 expression.

To develop appropriate preclinical models, we used p53^-/-^ mice as a sensitized background to generate Plk2^-/-^; p53^-/-^ murine mammary tumors in a manner similar to that used previously to develop a bank of transplantable p53^-/-^ tumors in Balb/c mice (45) (Supplemental Figure S4), the latter of which have been extensively characterized in our laboratories (46). Following transplantation of the Plk2^-/-^; p53^-/-^ MECs into the cleared fat pad of wild-type Balb/c recipients, palpable tumors were observed following an eight-month to greater than a one-year latency period, similar to that observed previously for p53^-/-^ MECs. RNA-seq analysis showed that they can be classified into three distinct molecular TNBC subtypes designated as luminal-like, basal-like, and claudin-low similar to p53^-/-^ mammary tumors (Supplemental Figure S5). Evidence for these classifications comes through the examination of overall expression patterns (Supplemental Figure S5) as well as known key subtype defining genes (Supplemental Figure S6).

To determine proper controls for our Plk2^-/-^; p53^-/-^ mammary tumors, we conducted a hierarchical clustering analysis on RNA-seq datasets of 192 murine mammary tumor samples, and identified the closest p53^-/-^ tumors for each Plk2^-/-^; p53^-/-^ TNBC subtype (Supplemental Table S2, Supplemental Figure S5). We next performed mammary fat pad transplantation for all the matched genotypes/subtypes of tumors and tested four treatment conditions: vehicle control, carboplatin alone, PLK1 inhibitor volasertib alone, as well as a combination of carboplatin and volasertib. Pilot experiments using volasertib [50 mg/kg/wk, a tolerated dose using colon carcinoma PDX models (47)] plus carboplatin showed a strong therapeutic effect in Plk2^-/-^; p53^-/-^ claudin-low tumors, but not p53^-/-^ claudin-low tumors (Supplemental Figure S7A and S7B). However, this dosage is also toxic as the mice had body weight loss and some mice in volasertib treatment groups, and carboplatin plus volasertib groups, eventually died (Supplemental Figure S7C and S7D). We then reduced the dose to 25 mg/kg/wk, which was well tolerated by all GEM models (Supplemental Figure S8). We found Plk2^-/-^; p53^-/-^ claudin-low, basal-like and luminal-like tumors showed a markedly better response to the combination treatment as compared to their paired p53^-/-^ tumors with wild-type Plk2 expression (Figure 4). Statistical analysis comparing different tumor types when the control treatment group reached ∼1500 mm^3^ confirmed the significant differences among these models as a function of Plk2 expression (Figure 4C, 4F, and 4I). Collectively, these data suggested that PLK1 inhibition in combination with chemotherapy is more effective in treating TNBC that has low PLK2 expression.

**Figure 4.**
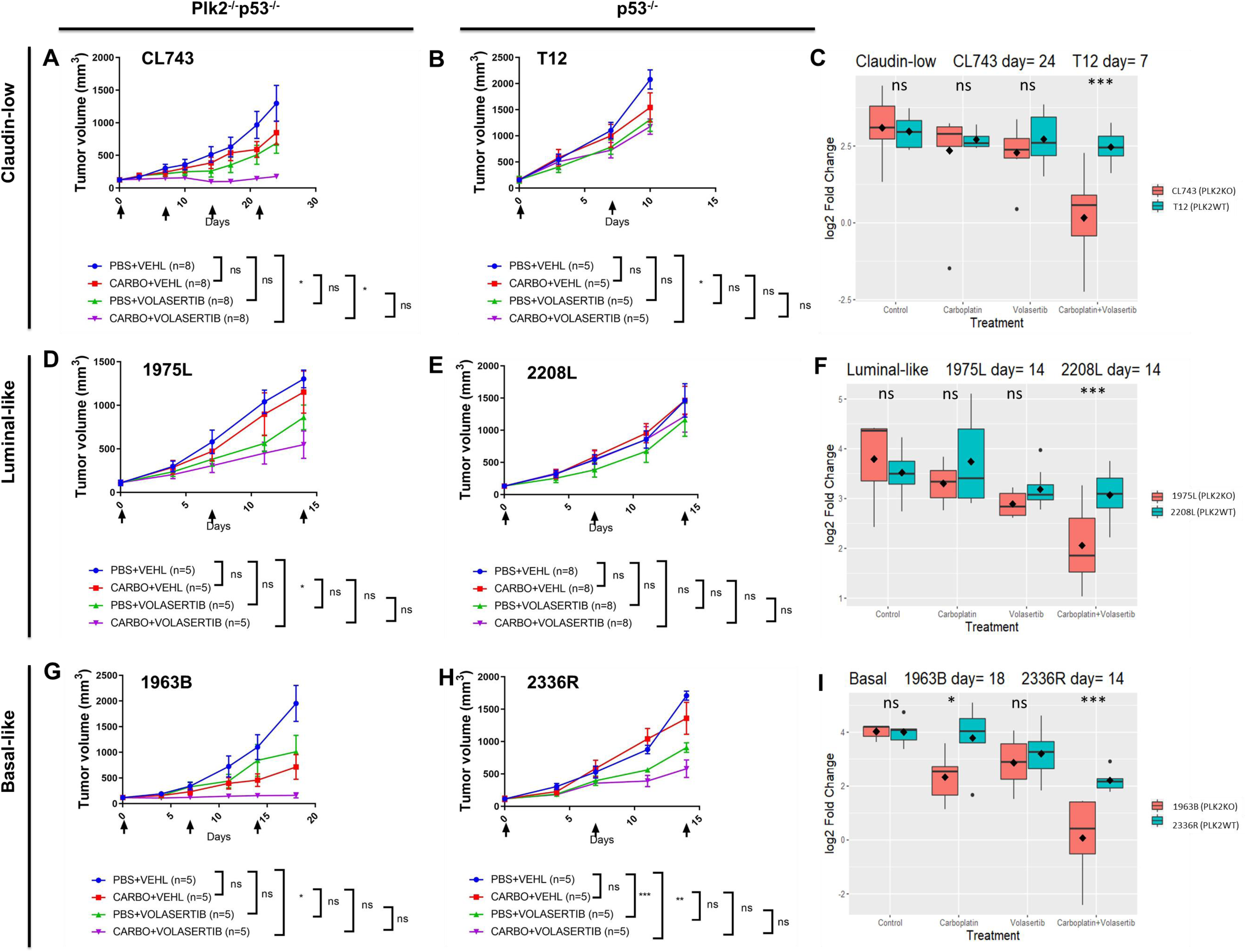
Plk2^-/-^; p53^-/-^ mammary tumors respond better to treatment with carboplatin plus the Plk1 inhibitor volasertib than their subtype-matched Plk2^+/+^; p53^-/-^ control tumors. Tumor growth curves of claudin-low subtypes CL743 and T12 **(A and B)**, luminal-like subtypes 1975L and 2208L **(D and E)**, basal-like subtypes 1963B and 2336R **(G and H)**. Tumor size was measured using a digital caliper twice a week until control groups reached ∼1500 mm^3^. Two-way ANOVA followed by Tukey test for multiple comparisons was used to analyze the growth curves of A, B, D, E, G, and H. Statistical comparison of the experimental endpoint was shown. **(C, F, and I)** Comparison of log2 fold change of tumor size between same treatment groups of Plk2^-/-^; p53^-/-^ tumors and p53^-/-^ tumors. Linear hypothesis test from the R package was used to compare the same treatment groups between different tumor models of C, F, I. Arrows indicate when the mice were treated. VEHL: vehicle; CARBO: carboplatin; ns, not significant; *, p <0.05; **, p < 0.01; ***p < 0.001.

In order to evaluate the clinical relevance of PLK2-PLK1 interaction in human tumors, we extended our study to several TNBC PDX models with high or low PLK2 expression (Supplemental Figure S9A) (https://pdxportal.research.bcm.edu/). We identified six PDX models from the Baylor College of Medicine (BCM) breast cancer PDX collection maintained by the Patient-derived Xenograft and Advanced In Vivo Models (PDX-AIM) Core Facility according to their PLK2 mRNA expression and known response to carboplatin from an ongoing preclinical trial, and tested the efficacy of volasertib in combination with carboplatin. We divided the six PDX models into three groups based on their response to carboplatin: complete responders, partial responders, and nonresponders. In both complete responders and partial responders to carboplatin, PDX models with lower PLK2 expression had a better response to volasertib, which induced tumor cytostasis (BCM-0002 vs. BCM-7482, BCM-2665 vs. BCM-15003) (Figure 5A, 5C, 5E, 5G, and 5I, Supplemental Figure S10 and S11). In contrast, tumor models with higher PLK2 expression continued to grow under volasertib administration, although at a slower rate compared to their controls (Figure 5B, 5D, 5F, 5H, and 5I, Supplemental Figure S10 and S11). Comparison of PLK2-low PDX lines to PLK2-high PDX lines when log2 fold change of control groups reached 2.5 suggested volasertib has a better tumor growth inhibitory effect in PLK2-low PDX lines (Figure 5E to 5I). However, in nonresponders, although volasertib slowed tumor growth of the model with lower PLK2 expression (BCM-4664), there was one exception where it induced a complete response in the PDX model with higher PLK2 expression (HCI-027) (Supplemental Figure S10).

**Figure 5.**
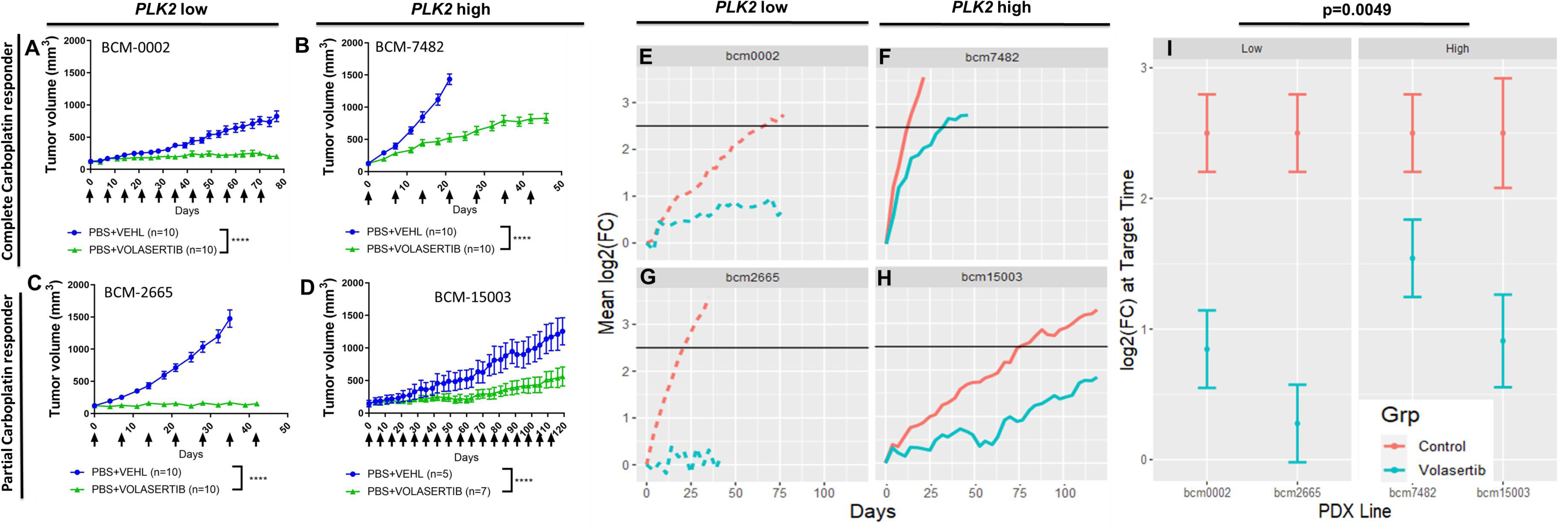
TNBC PDX with low *PLK2* expression had a better response to volasertib treatment than TNBC PDX with high *PLK2* expression if they showed response to carboplatin treatment. **(A and B)** BCM-0002 and BCM-7483 were complete responders to carboplatin treatment. **(C and D)** BCM-2665 and BCM-15003 showed partial response to carboplatin treatment. Note that volasertib treatment induced tumor cytostasis in BCM-0002 and BCM-2665. Tumor size was measured using a digital caliper twice a week. Two-way ANOVA followed by Tukey test for multiple comparisons was used to analyze the growth curves for each model. Statistical comparison at the endpoint of control group was shown. VEHL: vehicle; ****, p < 0.0001. **(E to H)** Mean log2 fold change (FC) from baseline for each PDX line and treatment group. The target log2(FC) (2.5 in the control group, black line) was used for statistical analysis of panel I. **(I)** A linear contrast was constructed to test whether the average difference in log2(FC) between control and volasertib in the PLK2 low PDX lines is the same as the average difference between control and volasertib in the PLK2 high PDX lines.

To understand the relationship between PLK2 loss and carboplatin treatment in predicting response to PLK1 inhibitors, we performed data mining on current clinical trials. Unfortunately, no clinical trial data are available using volasertib in combination with chemotherapy in TNBC patients. Nonetheless, by analyzing the recently published BrighTNess phase 3 neoadjuvant clinical trial in which carboplatin was added to standard neoadjuvant chemotherapy for TNBC patients (48), we observed a negative correlation between PLK1 and PLK2 expression in those patients whose tumors achieved a pathological complete response as compared to those with residual disease (Supplemental Figure S12). This provides additional support for the potential clinical relevance of PLK2 loss and carboplatin response.

To further determine the role of PLK2 in predicting the response to PLK1 inhibitor, we established two doxycycline-inducible isogenic models. First, we generated a mouse tumor cell line taking advantage of the residual neomycin cassette from the 1963B tumors (Plk2^-/-^; p53^-/-^ basal-like TNBC). Second, we examined the CNVs of the breast cancer cell lines from the Broad Institute. Surprisingly, almost none of the existing established human breast cancer cell lines exhibited a loss of PLK2 (see Discussion). Only the BT20 cell line was identified to exhibit partial PLK2 and MAP3K1 (another chromosome 5q marker) loss (Supplemental Figure S13). Interestingly, BT20 xenografts exhibited an extremely long latency in NSG mice (Supplemental Figure S14). We then infected both cell lines with the doxycycline-inducible vector pCW57.1 containing PLK2 and implanted the mouse 1963B-iPLK2 and human BT20-iPLK2 cells into the 4^th^ mammary fat pad of Balb/c and NSG mice, respectively. Restored PLK2 expression was validated by qPCR and western blot (Figure 6A and 6B, Supplemental Figure S14A and S14B). Volasertib treatment significantly reduced the tumor growth rate in both 1963B-iPLK2 and BT20-iPLK2 tumors in the absence of doxycycline induction (Figure 6C and Supplemental Figure S14C). Strikingly, doxycycline (DOX) induction of PLK2 expression resulted in decreased volasertib inhibition of tumor growth at the experimental endpoints in both models (Figure 6C, Supplemental Figure S14C and S15). Comparison of log2 fold change of tumor volume starting from the drug treatment to the experimental endpoint confirmed the reduced effect of volasertib in DOX treated 1963B-iPLK2 tumors (Figure 6D). Taken together, these studies provide support in isogenic models that PLK2 loss may serve as a potential biomarker to identify TNBC patients likely to be responsive to PLK1 inhibition.

**Figure 6.**
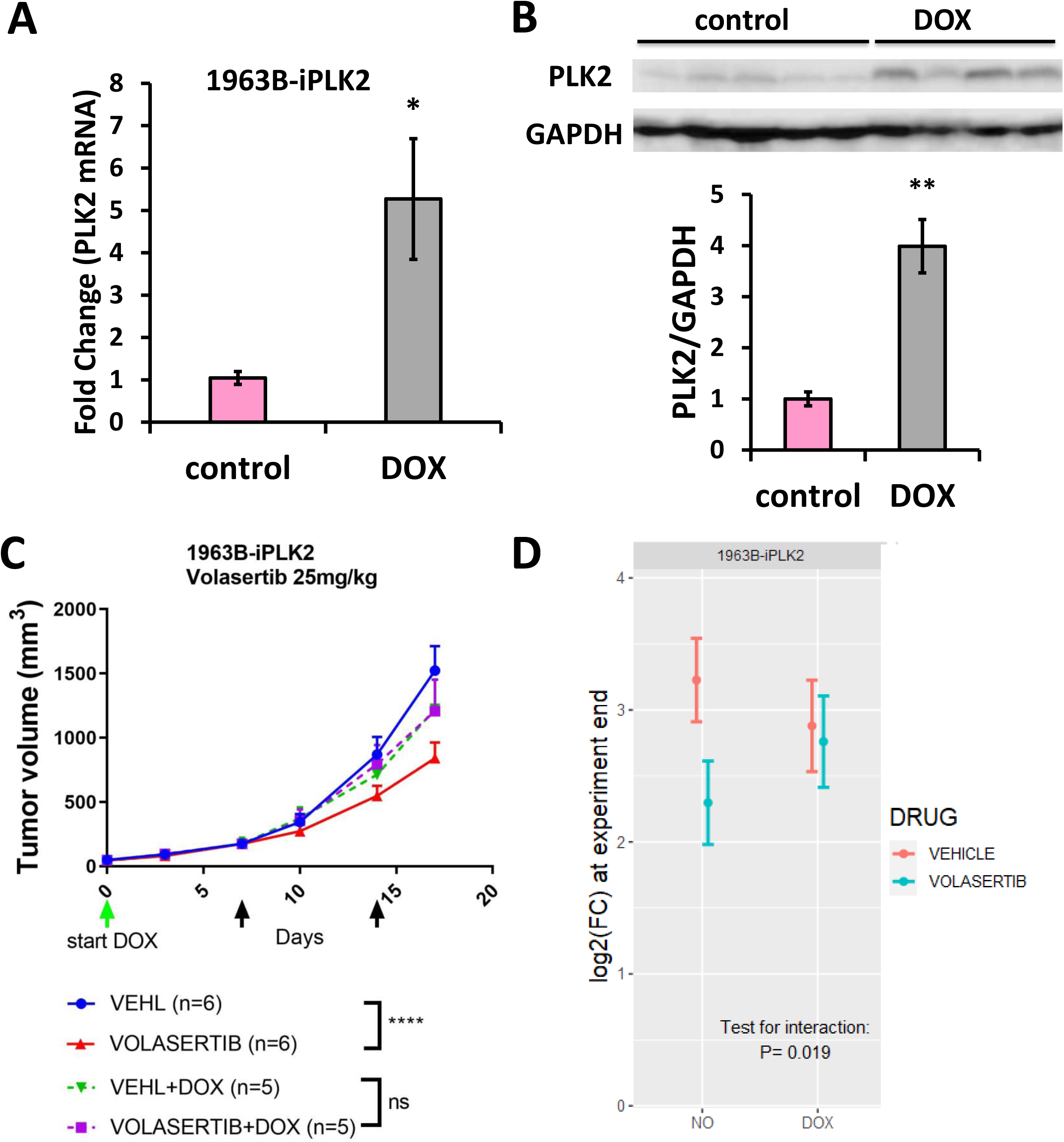
Doxycycline-induced PLK2 expression abolished the therapeutic effect of volasertib in mouse Plk2^-/-^; p53^-/-^ 1963B basal-like breast tumors. **(A)** qPCR analysis showed that PLK2 mRNA expression was induced by doxycycline water (DOX) in the mouse 1963B-iPLK2 tumors. n = 6 for 1963B-iPLK2 control group. n =5 for 1963B-iPLK2 DOX group. Statistical significance was determined by unpaired student’s t-test. *, p < 0.05. (B) Western blot analysis and quantification showed that PLK2 protein expression was induced by DOX in the mouse 1963B-iPLK2 tumors. n = 5 for 1963B-iPLK2 control group. n =4 for 1963B-iPLK2 DOX group. Statistical significance was determined by unpaired student’s t-test. **, p < 0.01. (C) Tumor growth curves of mouse tumors under control and volasertib treatment with or without DOX. Note that volasertib treatment significantly reduced tumor growth for non-DOX treated tumors but not DOX treated tumors at the experimental endpoints. Tumor size was measured using a digital caliper twice a week. Two-way ANOVA followed by Tukey test for multiple comparisons was used to analyze the growth curves. Statistical comparison of the experimental endpoint was shown. Green arrow indicates when DOX treatment began. Black arrows indicate when the mice were treated with drugs. VEHL: vehicle; ns, not significant; ****, p < 0.0001. (D) Two-way ANOVA analysis of log2 fold change (FC) of tumor volume starting from drug treatment to the end of experiment confirmed the reduced tumor inhibitory effect of volasertib in DOX treated 1963B-iPLK2 tumors.

## Discussion

Several chemotherapy drugs, either singly or in combination, including doxorubicin, cyclophosphamide, paclitaxel, and carboplatin, are still the standard-of-care therapy for many types of cancer, including breast cancer (49). However, these drugs have substantial side effects and limitations such as toxicity, immunosuppression, and drug resistance (50). In recent years, there has been a focus on developing new selective anti-mitotic drugs to help overcome some of these issues. PLK1 is a popular target as it regulates multiple essential steps of mitosis (51). Overexpression of PLK1 has been found in multiple types of solid tumors as well as in leukemia, and correlated with poor prognosis and survival (51). Several PLK1 inhibitors are currently in clinical development, including volasertib (52). In the current study, we found that PLK2 loss may provide a biomarker to guide PLK1 inhibitor patient selection, and furthermore, that it may be effective at a lower dose with reduced toxicity.

Unlike the well-established oncogenic function of PLK1, the role of PLK2 in human cancers remains unclear. PLK2 has been reported to be tumor suppressive in B-cell neoplasia, laryngeal carcinoma, cervical cancer cell lines and patient samples by promoting apoptosis and inhibiting cell proliferation (32,53,54). On the other hand, PLK2 expression was also correlated with improved cell survival using knockdown techniques in head and neck cancer and non-small cell lung cancer cell lines (41, 55). Contradictory effects of PLK2 in colorectal cancer also have been reported. One study showed that PLK2 promoted colorectal cancer growth and inhibited apoptosis by targeting Fbxw7/Cyclin E (56). In contrast, another study suggested that PLK2 was a tumor suppressor and its loss was more common in colorectal carcinoma than adenomas (57). Therefore, the role of PLK2 appears to be tumor type- and/or stage-dependent. In the present study, using a genetic shRNA screen, we observed that loss of PLK2 promoted colony formation of human mammary epithelial cells. Bioinformatic analysis of several human breast tumor datasets also indicated that PLK2 (chromosome 5q11.2) is located within a peak region of frequent deletion in breast cancer, consistent with previously published studies (35); moreover, the same chromosomal region is frequently deleted in several other cancers (9, 10). Furthermore, PLK2 mRNA expression is significantly lower in breast cancer as reported in the TCGA dataset. Therefore, it is conceivable that PLK2 is a tumor suppressor in breast cancer, specifically that chromosomal loss of PLK2 is highly enriched within the basal-like/triple-negative subtypes of breast cancer. In agreement with this finding, we found that PLK2 mRNA expression in TNBC is the lowest as compared to other breast cancer subtypes.

Despite the fact that Plk1 heterozygotes (Plk1^+/-^) and Plk2 null (Plk2^-/-^) mice being viable (27, 40), we observed a synthetic lethal phenotype of Plk2^-/-^; Plk1^+/-^ offspring, suggesting a potential genetic interaction between PLK2 and PLK1. Therefore, we tested whether PLK2 can directly interact with PLK1 using two independent methods: BiFC and PLA. Our efforts to identify the PLK2 interactome by mass spectrometry failed to identify PLK1 as a PLK2 substrate (unpublished data), most likely explained by the transient interaction of PLK1 and PLK2 during prometaphase. Two PLK2 mutants analyzed using the PLA revealed that the kinase domain, but not the polo-box domains, of PLK2 is responsible for their interaction at prometaphase. While these studies have helped shed light on interactions with target substrates and the importance of proper subcellular localization and cell cycle, the precise mechanism of how PLK2 might regulate PLK1 during prometaphase is still unclear. It has been reported that PLK2 can phosphorylate Ser-137 of PLK1 to promote human colon cancer cell survival in cells with mitochondrial dysfunction (58). Given that PLK2-PLK1 interaction at prometaphase was mediated by PLK2 kinase domain, it is tantalizing to speculate that PLK2 may phosphorylate PLK1 to regulate cell cycle progression in normal mammalian epithelial cells and that loss of PLK2 will change PLK1 subcellular location and cell cycle dependent activity, thereby leading to a mitotic catastrophe and promoting tumor growth. However, phosphorylation of Thr-210 in the T-loop of the PLK1 kinase domain by Bora/Aurora-A-dependent phosphorylation is thought to be required for mitotic entry (59). Thus, the underlying mechanism of how PLK2 regulates PLK1 warrants further investigation.

Despite the rapid progress in the development of PLK1 inhibitors, there is a lack of biomarkers that can predict a patient’s response to this treatment. A recent study showed that targeting PLK1 using volasertib induced synthetic lethality in BRCA1-deficient cells (60), thus suggesting BRCA1 may serve as a biomarker for the response to PLK1 inhibition treatment. Our findings that PLK1 might mediate PLK2 function both *in vitro* and *in vivo* encouraged us to investigate the implication of PLK2 loss in the context of PLK1 inhibitor treatment.

Volasertib has been reported to synergistically enhance the efficacy of radiation treatment both *in vitro* and *in vivo* in glioblastoma (61). PLK1 inhibition in combination with doxorubicin plus cyclophosphamide, or paclitaxel, also achieved a better outcome in the treatment of TNBC xenograft models (38, 62). However, in the latter studies, the level of PLK2 expression was not reported. We found that volasertib treatment sensitized PLK2-low TNBC tumors to carboplatin in our rapidly-growing GEM models across all of the three subtypes of TNBC studied. This was also observed in several pairs of PLK2 high and low PDX models with much more complex genetics, with one notable exception (HCI-027) where a marked response to volasertib was observed in the absence of low Plk2 expression; this may reflect the high level of PLK1 protein expression in this model (Supplemental Figure S9C), but further studies will be required to understand the mechanisms responsible. Thus, as expected from the complex genetics of TNBC, decreased PLK2 expression alone may not always suffice to predict response to PLK1 inhibitors, however, low PLK2 expression does appear to at least highly enrich for PLK1 inhibitor responsiveness. Support for this comes from our GEM and PDX models demonstrating that volasertib worked better in PLK2 low models when treated with carboplatin. Carboplatin is known to block DNA replication and transcription and induce cell death (63). However, the molecular mechanisms underlying the resistance and sensitivity of tumors to carboplatin are still under investigation. Given the important role of the PLK family in cell cycle and proliferation, tumors sensitive to carboplatin and volasertib may share some common mechanisms related to cell proliferation and cell death. Analysis of BrighTNess clinical trial which added carboplatin to standard neoadjuvant chemotherapy for TNBC patients supports the potential clinical relevance of PLK2 loss and carboplatin response. In agreement with this analysis, our study of restoring PLK2 in the mouse Plk2^-/-^; p53^-/-^ 1963B-iPLK2 basal-like tumors showed that carboplatin response was blocked by DOX-induced PLK2 expression (Supplemental Figure S15). Nonetheless, further studies are needed to uncover the underlying mechanisms.

The synthetic lethality observed between Plk1^+/-^ and Plk2^-/-^ mice indicates that PLK2 loss may sensitize tumors to lower doses of PLK1 inhibitors. In this study, treatment with carboplatin and volasertib was performed at approximately 50% of the reported clinically relevant doses of each as a single agent, without any apparent toxicity as revealed by weight loss or mortality. These results suggest that it may be possible to reduce the toxicity of these individual agents by dose de-escalation studies done in combination, though this issue is beyond the scope of this study or the statistical power of the mouse studies performed. Also, as the combinational drug treatment regimens evolving (64), it will be interesting to examine whether carboplatin and volasertib can be administrated in a sequential treatment strategy or an alternating dosing schedule.

To identify a human breast cancer cell line with PLK2 loss for our study, we screened the Broad Institute breast cancer cell line database and surprisingly found that the deletion of PLK2 and other nearby chromosome 5q genes was rarely observed in breast cancer cell lines in contrast to established PDX models and patient TCGA datasets (Figure 1, Supplemental Figure S1 and S13). This suggests that loss of PLK2/ chromosome 5q was selected against under *in vitro* cell culture conditions. Furthermore, PDX models are a better representation of the tumor status in patients. Nevertheless, we identified BT20 as a PLK2 low model and generated a doxycycline-inducible PLK2 system. In previous studies, both when implanted subcutaneously or orthotopically in the mammary fat pad, BT20 cells exhibit a delayed growth response and have recently been shown to fail to metastasize following intracardiac injection (65–68). Notwithstanding the delayed growth kinetics, together with the doxycycline-inducible PLK2 mouse tumor model, we demonstrated that a higher level of PLK2 expression was able to impair the therapeutic effect of volasertib at the experimental endpoint. Thus, these results obtained in both mouse and human isogenic models help support the role of PLK2 as a biomarker for predicting response to the PLK1 inhibitor.

Volasertib is an ATP-competitive compound for the PLK1 kinase domain and we found that the kinase domain of PLK2, but not the PBD, played an essential role in PLK2-PLK1 interaction. It will be interesting to examine whether TNBC patients with low-PLK2 expression can benefit from the treatment with PBD-binding antagonists, another class of PLK1 inhibition drugs. Since PBD is unique for the PLK family and required for their functions, compounds specially targeting PBD represent an ideal group of PLK inhibitors. The first PBD inhibitor poloxin was shown to inhibit breast cancer xenograft growth by suppressing proliferation and triggering apoptosis (69, 70). However, it was reported as a non-specific protein alkylator in a recent study (71). The natural benzotropolone compound purpurogallin (PPG) was identified as a PLK1 PBD inhibitor in an *in vitro* screen, but later was found to interfere with various other proteins due to its structural features (71, 72). Poloppins, however, seemed to be more specific PBD inhibitors and were shown to effectively target KRAS-expressing colorectal cancer xenografts (73). Therefore, comparing poloppins to volasertib may yield more information about PLK2-PLK1 interactions and benefit breast cancer patients with a more precise treatment.

Beroukhim et al. reported previously that chromosome 5q loss containing PLK2 was one of the top 20 most significant peak deletion regions detected across 26 types of human cancers (9). Taylor et al (10) and our unpublished studies (Siegel, Perou, et al., manuscript in preparation), have also revealed that chromosome 5q loss often occurs in lung squamous cell carcinoma, ovarian cancer, as well as TNBC. Therefore, the utility of PLK2 as a biomarker may have a broader application and provide a therapeutic window for the use of PLK1 inhibitors in multiple types of cancers.

## Materials and Methods

### Anchorage-independent proliferation assay

TLM-HMECs [genetically engineered human mammary epithelial cells, (36)] were lentivirally transduced with control or human PLK2 GIPZ lentiviral shRNAs (Open Biosystems-Horizon Discovery, Supplemental Table S3), and then selected using puromycin to establish stable cell lines. These cell lines were subsequently transduced with lentiviruses expressing control or human PLK1 GIPZ lentiviral shRNA (Open Biosystems-Horizon Discovery, Supplemental Table S3), at an MOI of 1 and subjected to anchorage-independent proliferation assay as described (37). Cells were seeded at densities of 3 × 10^4^ or 5 × 10^4^ per 60 mm plate with a bottom layer of 0.6% Noble agar in MEM (Gibco) and a top layer of 0.5% methylcellulose containing MEGM (Lonza). Fresh MEGM (0.5 ml) was added every three days. Macroscopic colonies were counted after four weeks. Experiments were performed in triplicates.

### Primary mammary epithelial cell isolation, transduction, and transplantation

Primary mammary epithelial cells (MECs) were isolated from 8-week-old Plk2^-/-^ mice for transplantation experiments. MECs were isolated by mincing freshly harvested mammary glands into 1 mm^3^ fragments using a Vibratome Series 800-Mcllwain Tissue Chopper. The tissue fragments were digested in DMEM/F12, which contained 2 mg/ml collagenase A (Roche Applied Science), for 1 hr at 37°C shaking at 120 rpm. The adipocytes were removed from the organoids by centrifugation at 1500 rpm for 5 min. Following this centrifugation step, the remaining stromal cells were removed by sequential centrifugation at 1500 rpm for 5 sec. The organoids were then subjected to trypsinization by resuspending them in 0.25% Trypsin-EDTA for 5 min at 37°C and subsequently the organoids were washed and filtered using a 40 µm cell strainer to obtain a single-cell suspension.

A LeGO-T lentiviral vector was kindly provided by Kristoffer Riecken (74). shRNAs targeting mouse Plk1 (Supplemental Table S3) were purchased from Open Biosystems-Horizon Discovery. U6 promoter-hairpin segments were amplified from a pLKO.1 vector using PCR primers XbaI F: gagatctagaccttcaccgagggcctatttc and NotI R: gagagcggccgcccatttgtctcgaggtcgag. Both LeGO-T lentiviral vector and PCR fragments were digested with XbaI + NotI, purified with QIAquick Gel Extraction kit or QIAquick PCR Purification kit (Qiagen), ligated using T4 DNA Ligase (New England BioLabs), and transformed into One Shot Stbl3 competent cells (Life Technologies). All plasmids were sequence-verified prior to lentiviral production. Lentiviral vectors were co-transfected with packaging vectors VSVG and gag/pol in a ratio of 3:1:2 into HEK293T cells using Trans-IT Transfection reagent (Mirus). Viral supernatants were collected at 48 and 72 hr post-transfection, pooled, and filtered through 0.45 µM filters to remove cellular debris. Filtered viral supernatants were concentrated using Beckman Coulter Optima ultracentrifuge (SW32Ti rotor) at 25,000 rpm for 1 hr 45 min. Ultracentrifuged viruses were resuspended in MEC growth media and titered by FACS analysis as described before (75).

Primary MECs were plated in a non-adherent dish and were transduced with the above lentiviral Plk1 at an MOI of 50. Cells were incubated at 37°C overnight. The next day cells were washed to remove any unbound virus and were resuspended in DMEM/F12 containing 20% Matrigel at a concentration of 20,000 cells/µl and kept on ice until transplantation. For transplants, 150,000 cells were injected into cleared contralateral fat pads of 3-week-old SCID/Beige host mice. Mammary glands were harvested 8 weeks post-transplantation and whole-mount, as well as histological analyses, were performed.

### Estrogen and progesterone treatment

To induce proliferation in mammary epithelial cells and quantify the mitotic spindle orientation, mice were treated with estrogen and progesterone for two days as described before (29). In brief, mice were injected with 100 μl of estrogen and progesterone sesame oil solution with a final concentration of 1 μg of E2 and 1 mg of progesterone under the skin between the shoulder blades.

### Tissue harvest

The fourth pair of mammary glands was harvested at 8 weeks post-transplantation. BrdU (B5002, Sigma-Aldrich) at 60 μg/g body weight was i.p. injected 2 hr prior to tissue harvest, which allowed for proliferation analysis.

### RNA isolation

The fourth pair of mammary glands was isolated and MECs were purified from these glands. Total RNA was isolated using Trizol Reagent (Invitrogen) or RNesay Mini Kit (Qiagen) according to the manufacturer’s protocols. Similar RNA preparations were also done for the tumor samples.

### Protein extraction and immunoblot analysis

Protein was isolated as described before (69). Protein was quantified using a Pierce BCA Protein Assay Kit (Thermo Fisher Scientific). SDS-PAGE was employed to separate proteins that were transferred onto a PVDF membrane. PLK1 (Abcam, ab17057, 1:1000), PLK2 (Cell Signaling Technology, 14821, 1:1000), and GAPDH (Cell Signaling Technology, 2118, 1:5000) antibodies were employed.

### Whole-mount, carmine alum staining, and branching analysis

For mammary gland whole-mount analysis, tissue was mounted between two glass slides and imaged using Leica MZ16F fluorescence stereoscope. Images (1.0x and 1.6x) of control luciferase and Plk1 shRNA lentiviral vector-transduced mammary glands were taken for branching analysis. Following these analyses, mammary glands were paraffin-embedded. Carmine alum staining was performed as described before (29).

### Immunofluorescence

Paraffin-embedded tissues were cut into 5 μm sections and dried before use. To begin, sections were deparaffinized in xylene and rehydrated in graded ethanol solutions. Antigen retrieval was performed by boiling in 10 nM sodium citrate buffer (pH 6.0) for 20 min. Washing steps were performed with 1×PBS and primary antibodies were incubated at 4°C overnight in a humidified chamber. All primary antibodies (BrdU, Abcam, ab6326, 1:1000; NuMA, Abcam, ab36999, 1:250) were diluted in 5% BSA, 0.5% Tween-20 blocking buffer.

### Bimolecular fluorescence complementation assay

pDONR vectors encoding PLK2, RB1, CHK1, TUBB, and PLK1 were obtained from human ORFeome (Open Biosystems-Horizon Discovery). They were individually transferred into either bait pB-CMV-CVn-neo (for PLK2) or prey pB-CMV-YFP-CC-puro (for RB1, TUBB, CHK1, PLK1) vectors using Gateway recombination reaction (Life Technologies) (76). Bait and prey plasmids containing fused fragments were then individually transfected together with VSVG and gag/pol packaging vectors in a ratio of 3:1:1 into HEK293T cells using Trans-IT Transfection reagent (Mirus) to produce retroviruses. Stable cell lines were generated after selection with G418 (500ug/ml) (bait) and puromycin (2ug/ml) (prey), and the expression of proteins was verified by immunofluorescence and western blotting. In BiFC assay, the PLK2-bait cell line was infected with one of the CHEK1, TUBB, RB1, or PLK1 prey-produced retroviruses.

Fluorescent signals were observed after 48 hr of infection using a Carl Zeiss inverted fluorescence microscope with an AxioCam MRm camera. BD LSRII Flow Cytometer (BD Biosciences) was utilized to quantify YFP-positive cells. Untransduced cells, cells transduced with bait-empty vector (without PLK2 fusion), or cells transduced with only one expression vector containing a fused fragment (either bait or prey) were used as negative controls. BiFC assays were repeated five times in triplicates.

### Proximity ligation assay

Full-length PLK1 and PLK2 sequences were PCR amplified and individually cloned into pCDNA3.1 vector (Invitrogen) using XbaI-EcoRI and XbaI-BamHI (New England BioLabs) restriction sites, respectively. PLK2 mutations K111R (KD, kinase dead), W503F (PBD, polo-box domain 1) and H629A, K631M (PBD, polo-box domain 2) were generated by site-directed mutagenesis (Agilent Technologies) according to the manufacturer’s protocol. All constructs were verified by DNA sequence analysis prior to transfections.

HEK293T cells were maintained in Dulbecco’s modified Eagle medium (DMEM) supplemented with 10% fetal bovine serum (GenDepot) and antibiotic-antimycotic solution (GenDepot). Cells were seeded at 40% confluence before the day of transfection. To induce an arrest in the G2/M phase of the cell cycle, cells were treated with 40 ng/mL of nocodazole (Sigma-Aldrich) for 16 hrs. For each transfection, 2.5 ug of total DNA was mixed with 250 ul of Opti-MEM I Reduced-Serum medium (Life Technologies) and 7.5 ul of Trans-IT-293 Reagent (Mirus). The mixture was incubated at room temperature for 20 min and then added to cells. After 48 hr cells were transferred into a 96-well SensoPlate with a glass bottom (Thermo Scientific) in the amount of 10^4^ cells per well and were grown for another 24 hr. Subsequently, cells were fixed with 4% paraformaldehyde (Thermo Scientific) for 10 min, permeabilized with 0.25% TritonX-100 for 10 min, and blocked with 3% BSA in PBS for 1 hr at room temperature. Then, cells were co-incubated with mouse anti-PLK1 antibody (Abcam, ab17057) diluted 1:4000 and rabbit anti-PLK2 antibody (Cell Signaling, 14812) diluted 1:2000 overnight at 4°C. The PLA was performed using reagents supplied in the Duolink in situ Red PLA Mouse/Rabbit kit (Sigma-Aldrich, DUO92101) following the manufacturer’s instructions. Cell nuclei were stained with NucBlue ReadyProbes reagent (Invitrogen), and actin filaments were stained with Actin Green 488 ReadyProbes reagent (Invitrogen).

Imaging was performed on a Cytiva DV Live epifluorescence image restoration microscope using an Olympus PlanApo N 60x/1.42 NA objective and 1.9k x 1.9k pco.EDGE sCMOS_5.5 camera with 1042×1042 FOV. The filter sets used were DAPI (390/18 excitation, 435/48 emission) and CY5 (632/22 excitation, 676/34 emission). Z-stacks (0.25 μm) covering the whole cell (∼8.3 μm) were acquired before applying a conservative restorative algorithm for quantitative image deconvolution using SoftWorx v7.0 and saving files in pixel intensity projection tiff format for each channel. Imaging for quantitation of PLA positive cells was done on a BioTek Cytation 5 Cell Imaging Multi-Mode Reader equipped with a DAPI filter cube set (excitation 377/50, emission 447/60) and CY5 filter cube set (excitation 628/40, emission 685/40) with a Grasshopper3 GS3-U3-14S5M camera. Image panels were collected with an Olympus 20x/0.45NA objective. Images were exported from the Gen5 version 3.03.10 software as greyscale tiff files. Quantitative PLA analysis was performed by calculating the percentage of cells showing positive PLA signals from eight randomly captured image panels per group.

### RNA-seq and bioinformatic data analysis

RNAseq was performed as described before (77). Briefly, Paired-end (2×50bp) sequencing was performed on the Illumina HiSeq 2500 sequencer at the UNC High Throughput Sequencing Facility. RNA-seq results were aligned to the mouse mm10 reference genome using the Star alignment algorithm (78) and quantified as gene-level counts using a Salmon pipeline as previously described (77). Upper-quartile normalized counts were then log2(x+1) transformed. Hierarchical clustering was performed with 1910 intrinsic mouse genes using Cluster 3.0 with 1-Pearson correlation distance and centroid linkage (79).

To compare the gene expression of Plk2^-/-^; p53^-/-^ models with published mouse model gene expression classes derived from microarray expression, we combined the log2(x+1) RNA-seq expression data with the expression data from microarray samples previously used to identify mouse mammary tumor expression classes (79). Batch effects due to expression type (RNA-seq vs microarray) were adjusted using COMBAT (80). Expression class assignments from the initial publication (79) were retained for microarray samples. Plk2^-/-^; p53^-/-^ samples were assigned to three separate expression classes (PLK2-Luminal, PLK2-Basal, and PLK2-Claudin-low) based on the hierarchical clustering with RNA-seq samples. Data was uploaded to GEO (GSE174683).

PLK2 and PLK1 mRNA expression levels in PDX models were evaluated in the BCM PDX Portal (https://pdxportal.research.bcm.edu/).

Copy number data for TCGA breast cancer samples were downloaded from cBio portal (www.cbioportal.org) using the TCGA Firehose Legacy version. Copy number variation was calculated using GISTIC 2.0 with values < −2 indicating a deletion event – potentially homozygous deletion; values < −1 indicating a loss (potentially a heterozygous deletion), values near 0 indicating diploid status, values > 1 indicating a gain, and values >2 indicating an amplification.

### Generation of doxycycline-inducible cell lines

To derive a cell line from the 1963B Plk2^-/-^; p53^-/-^ tumor model, a fresh tumor was harvested when it reached ∼1cm in diameter. The tumor was then minced into small 1-2 mm^3^ pieces and digested with 1 mg/ml Type I Collagenase in DMEM/F12 for 2 hr in a 37 °C incubator shaking at 125 rpm. After centrifugation at 1500 rpm for 5 min, the pellet was resuspended in PBS and then three short centrifugations (1500 rpm at 7 seconds) were performed to enrich the mammary epithelial organoids. Single cells were obtained by digestion in 0.25% Trypsin-EDTA for 5 min at 37 °C. Since the p53 null mice have a neomycin resistance cassette, the cell line was generated using 500 µg/ml G418 (Sigma) selection for two weeks and then validated by pan-cytokeratin and cytokeratin 5 immunofluorescence staining.

Doxycycline-inducible vector pCW57.1 was obtained from Addgene (#41393). PLK2 in a pDONR223 vector was obtained from human ORFeome (Open Biosystems-Horizon Discovery) and was cloned into pCW57.1 using the Gateway recombination reaction (Life Technologies) and verified by sequencing prior to transfection. For viral production, the plasmid was transfected together with packaging plasmids CMV-VSVG, MDL-RRE, and RSV-REV in the ratio of (3:1:1:1) into HEK293T cells using Trans-IT transfection reagent (Mirus). Virus-rich supernatant was collected at 72 hr post-transfection, concentrated by ultracentrifugation at 25,000 rpm for 1 hr 45 min, and titered using lentivirus qPCR titer kit (ABM). 1963B and BT20 cells were transduced with the virus at an MOI of 10 following puromycin selection (2 ug/ml) for 5 days. All cell lines were cultured in DMEM/High Glucose medium (GenDepot) supplemented with 10% fetal bovine serum plus antibiotic-antimycotic solution (GenDepot) and grown in a humidified incubator at 37°C with 5% CO_2_.

### Mice, tumor models, and treatment

All animal experiments were conducted in accordance with a protocol approved by the Institutional Animal Care and Use Committee of Baylor College of Medicine and in compliance with all relevant ethical regulations regarding animal research. Balb/c mice were purchased from Envigo. NSG mice were purchased from the Jackson laboratory.

To generate Plk2^-/-^; p53^-/-^ mice, Plk2^-/-^ mice were first backcrossed with wild-type Balb/c mice for >5 generations and then bred with the p53^-/-^ Balb/c mice. To get breast-specific tumors, MECs isolated from Plk2^-/-^; p53^-/-^ mice were transplanted into cleared fat pads of wild-type Balb/c recipients. Spontaneous tumors arose with a latency of eight months to more than a year and were characterized by histology, H & E, immunofluorescence staining, and RNA-seq. Those tumors were cryopreserved as small chunks for later mammary fat pad implantation experiments as described before (81). PDX models were obtained from the Patient-Derived Xenograft and Advanced In Vivo Models Core at BCM.

When tumor size became ∼110-200 mm^3^, animals were randomized into different treatment groups. Carboplatin (Sigma-Aldrich, C2538) was reconstituted in PBS and delivered once a week at a dosage of 25 mg/kg or 50 mg/kg by i.p. injection. Volasertib (Selleckchem, S2235) was formulated in 0.1N hydrochloric acid (vehicle) and administered once a week by oral gavage at a dosage of 25 mg/kg or 50 mg/kg. Tumor size was measured using a digital caliper twice a week.

Doxycycline-treated mice were supplied with 200 µg/ml doxycycline water when their tumors became palpable. Doxycycline water was changed twice a week.

## Authors’ Contributions

Conception and design: Y. Gao, T.F. Westbrook, C.M. Perou, J.M. Rosen

Development of methodology: Y. Gao, E. Kabotyanski, E. Villegas, J. H. Shepherd, D. Acosta, T. Sun, C Montmeyor-Garcia, X. He

Acquisition of data (provided animals, acquired and managed patients, provided facilities, etc.): Y. Gao, E. Kabotyanski, E. Villegas, D. Acosta, C. Hamor, T. Sun, C Montmeyor-Garcia, L.E. Dobrolecki, M.T. Lewis

Analysis and interpretation of data (e.g., statistical analysis, biostatistics, computational analysis): Y. Gao, E. Kabotyanski, E. Villegas, J. H. Shepherd, T. Sun, X. He, S.G. Hilsenbeck, X.H.-F. Zhang

Writing, review, and/or revision of the manuscript: Y. Gao, E. Kabotyanski, E. Villegas, J. H. Shepherd, T.F. Westbrook, M.T. Lewis, X.H.-F. Zhang, C.M. Perou, J.M. Rosen

Administrative, technical, or material support (i.e., reporting or organizing data, constructing databases): Y. Gao. E. Kabotyanski, L.E. Dobrolecki, T.F. Westbrook, M.T. Lewis, X.H.-F. Zhang, C.M. Perou, J.M. Rosen

Study supervision: J. M. Rosen

## Acknowledgments

This study was funded by the following grants: Susan G. Komen (SAC110031, Rosen; SAC160074, Perou), NCI Breast SPORE program (P50-CA58223), and the NIH (R01-CA148761, Perou and Rosen). Imaging for this project was supported by the Integrated Microscopy Core at Baylor College of Medicine with funding from NIH (DK56338 and CA125123), CPRIT (RP150578, RP170719), the Dan L. Duncan Comprehensive Cancer Center, and the John S. Dunn Gulf Coast Consortium for Chemical Genomics. This project was also supported by the Cytometry and Cell Sorting Core at Baylor College of Medicine with funding from the CPRIT Core Facility Support Award (CPRIT-RP180672), the NIH (P30 CA125123 and S10 RR024574) and the expert assistance of Amanda White and Joel M. Sederstrom. We thank the Patient-Derived Xenograft and Advanced In Vivo Models Core at Baylor College of Medicine (Michael T. Lewis, Ph.D., Academic Director, Lacey E. Dobrolecki, M.S., Technical Director. Grants supporting the core: CPRIT Core Facility Award (RP170691) and P30 Cancer Center Support Grant (NCI-CA125123)). We thank the Biostatistics and Informatics Shared Resource Core (Susan G. Hilsenbeck, Ph.D., Director. Grants supporting: CCSG P30CA125123 and SPORE P50CA186784).

We thank Dr. Meenakshi Anurag for her help with the BrighTNess clinical trial analysis. We thank Dr. Chad A. Shaw for the initial bioinformatic analyses of TCGA data. We thank Ramakrishnan Rajaram Srinivasan for analyzing PLK2 and PLK1 mRNA expression of PDX models. We thank Hannah Johnson for assisting with imaging. We thank Shirley Small for her help with animal experiments. We thank Dr. Kristen L. Karlin for providing the human PLK1 shRNA. We thank Dr. Yating Cheng for her help with cloning. We thank Dr. Xi Chen for sharing the equipment. We thank Dr. Alana Welm, Dr. Helen Piwnica-Worms, and Dr. Gloria Echeverria for their helpful discussion.

