## Supplemental Figures, Tables, and Legends for "Tumor suppressor PLK2 may serve as a biomarker in triple-negative breast cancer for improved response to PLK1 therapeutics"

### Supplemental Figure Legends

**Supplemental Figure S1. PLK2 is tumor suppressive and knockdown of PLK1 rescues PLK2 loss induced transformation of HMECs. (A)** PLK2 is significantly deleted in 8 of 11 independent cancer types (q<=0.25) analyzed in the TCGA Pan-Cancer dataset (2013-08-16). Among these, PLK2 is located within a peak region of deletion in breast cancer and lung adenocarcinoma. PLK2 **(A)** and PLK1 **(B)** mRNA expression by ER/HER2 status. P-values were generated by pairwise T-tests with Bonferroni correction for multiple comparisons. **(D)** Knockdown of PLK2 in HMECs using two shRNAs, PLK2-296 and PLK2-309, both increased the colony number. This induction was abolished by additional knockdown of PLK1. Fifty thousand HMECs were seeded in the plates. Assays were performed in triplicate. **(E)** Knockdown efficiency of hPLK2 #1 and hPLK2 #2 shRNAs in HMECs. **(F)** Knockdown efficiency of hPLK1 shRNA in HMECs.

**Supplemental Figure S2. Plk1 shRNAs knockdown in primary MECs.** Primary MECs were isolated from $Plk2^{-/-}$ 8 wk old mice and transduced with lentiviruses. The shRNAs knockdown efficiency was evaluated by qPCR **(A)** and western blot **(B)**. mPlk1 #1 shRNA achieved 50% knockdown and mPlk1 #2 shRNA 75% knockdown of mRNA levels. The protein levels of PLK1 were reduced by 52% and 93% by mPlk1 #1 and mPlk1 #2 shRNA, respectively.

**Supplemental Figure S3. Proximity ligation assay of PLK2 and PLK1 after Nocodazole synchronization. (A)** Negative controls for PLA were performed in the absence of primary antibodies. No positive PLA signals were detected in untransfected cells (image a); in cells expressing only one protein: PLK1 (image b), PLK2 (image c), PLK2-PBD mutant (image e), or PLK2-KD mutant (image g); or in cells co-expressing PLK1 with PLK2 (image d), PLK1 with PLK2-PBD (image f), or PLK1 with PLK2-KD (image h) when only secondary antibodies have been used. **(B)** PLA in HEK293T cells co-expressing PLK1 and either PLK2 or PLK2 kinase-dead mutant (KD). Representative images for untreated cells (a-e) and cells treated with nocodazole (f-k) are shown. Positive PLA signals were obtained in cells co-expressing PLK1 with wild-type PLK2 (images c and h) or with PLK2-KD mutant (images e and k). Negative controls: PLA performed in non-transfected cells (images a and f); PLA performed in the absence of primary antibodies for cells co-expressing PLK1 with PLK2 (images b and g) and cells co-expressing PLK1 with PLK2-KD mutant (images d and j). Red: PLA signals. Blue: DAPI staining. Green: ß-actin staining. **(C)** Quantitative of PLA signals in cells co-expressing PLK1 with ether wild-type PLK2 or PLK2-KD mutant after treatment with nocodazole (40ng/ml) for 16 hr (G2/M arrest) using microscopy images. About one hundred PLA-positive cells were counted per group. Statistical significance was determined by unpaired student's t-test.

**Supplemental Figure S4. Generation of Plk2$^{-/-}$; p53$^{-/-}$ breast tumors. (A)** Schematic diagram of Plk2$^{-/-}$; p53$^{-/-}$ breast tumors generation. Briefly, p53$^{-/-}$ mice in Balb/c background were crossed with Plk2$^{-/-}$ mice in the same background to generate Plk2$^{-/-}$;

p53$^{-/-}$ mice. Then mammary epithelial cells were isolated from Plk2$^{-/-}$; p53$^{-/-}$ mice and implanted into wild-type Balb/c mice with cleared mammary fat pad to wait for tumor growth. **(B)** Primary Plk2$^{-/-}$; p53$^{-/-}$ breast tumors had a latency of 8 months to more than one year. We then collected the tumors and cryopreserved the tumor pieces for later implantation.

**Supplemental Figure S5. Hierarchical clustering of RNA-seq expression from Plk2$^{-/-}$; p53$^{-/-}$ tumors and other mouse mammary tumor RNAseq samples. (A)** Full hierarchical clustering of all samples by 1910 intrinsic mouse genes. Colored bars mark the location of Plk2$^{-/-}$; p53$^{-/-}$ tumor samples. **(B)** Clustering dendogram and colored bar indicating mouse model show Plk2$^{-/-}$; p53$^{-/-}$ tumors clustering on three distinct branches. The three groups of Plk2$^{-/-}$; p53$^{-/-}$ tumors are indicated by shaded bars. PLK2-Claudin-low samples (yellow) show low expression of Cldn3 and Cldn7. PLK2-Luminal tumors show low expression of basal markers and elevated expression of luminal gene clusters. PLK2-basal samples show high expression of the gene cluster including the Basal marker gene Krt5. Particular tumor lines used in this study are labeled and indicated by colored triangles above names. Plk2$^{+/+}$; p53$^{-/-}$ samples used as controls are shown with a black box around their name and a black outline for the triangle.

**Supplemental Figure S6. Plk2$^{-/-}$; p53$^{-/-}$ tumors exhibit characteristics of multiple gene expression classes.** Comparison of RNA expression from Plk2$^{-/-}$; p53$^{-/-}$ tumors and other mouse mammary tumor samples classified by expression subtype. (**A)** All subtypes of Plk2$^{-/-}$; p53$^{-/-}$ tumors have low Plk2 expression. **(B)** PLK2-Luminal subtype has high Met expression, similar to tumors of the p53-null-Luminal expression class. **(C)** PLK2-Basal subtype has elevated Krt5 expression. **(D)** PLK2-Claudin-low subtype has low expression of the Claudin gene Cldn7.

**Supplemental Figure S7.** $Plk2^{-/-}$; $p53^{-/-}$ claudin-low breast tumors had a better response to carboplatin plus PLK1 inhibitor volasertib treatment than $p53^{-/-}$ claudin-low tumors **(A and B)**. However, 50 mg/kg/wk of volasertib is toxic as the mice had body weight loss and some mice in volasertib treatment groups and carboplatin plus volasertib groups eventually died **(C and D)**. Tumor size was measured using a caliper twice a week until control groups reached ~1500 $mm^3$. Two-way ANOVA followed by Tukey test for multiple comparisons was used. Statistical comparison of the experimental endpoint was shown. VEHL: vehicle; CARBO: carboplatin; ns, not significant; *, $p < 0.05$; **, $p < 0.01$; ***, $p < 0.001$; ****, $p < 0.0001$.

**Supplemental Figure S8. Body weight of $Plk2^{-/-}$; $p53^{-/-}$ and $p53^{-/-}$ GEM models during treatment indicated 25 mg/kg/wk of volasertib was a tolerable dosage.** Two-way ANOVA followed by Tukey test for multiple comparisons was used to compare the different treatment groups. Statistical comparison of the experimental endpoint was shown. VEHL: vehicle; CARBO: carboplatin; ns, not significant. *, $p < 0.05$.

**Supplemental Figure S9. PLK2 and PLK1 expression in PDX models.** *PLK2* **(A)** *and PLK1* **(B)** *mRNA* expression levels for 90 breast cancer PDX models. Red indicated the

PLK2 high models while green indicated the PLK2 low models used in the treatment study. **(C)** PLK2 and PLK1 western blots of the 6 PDX models in the treatment study. n = 4 per model. Protein samples were run and transferred at the same time and signals were developed and captured at the same time for the two membranes.

**Supplemental Figure S10. TNBC PDX with low *PLK2* expression had a better response to volasertib treatment than TNBC PDX with high *PLK2* expression if they showed response to carboplatin treatment. (A and B)** BCM-0002 and BCM-7482 were complete responders to carboplatin treatment. **(C and D)** BCM-2665 and BCM-15003 showed partial response to carboplatin treatment. Note that combinational treatment of carboplatin and volasertib led to tumor regression in PLK2 low model BCM-2665 but not in PLK2 high model BCM-15003. **(E and F)** BCM-4664 and HCI-027, however, didn't respond to carboplatin treatment and had a different volasertib response pattern compared to other groups. Tumor size was measured using a caliper twice a week. Two-way ANOVA followed by Tukey test for multiple comparisons was used to analyze the growth curves for each model. Statistical comparison at the endpoint of control group was shown. VEHL: vehicle; ns, not significant; *, $p < 0.05$; **, $p < 0.01$; ***, $p < 0.001$; ****, $p < 0.0001$.

**Supplemental Figure S11. Body weight of all six PDX models during treatment indicated 25 mg/kg/wk of volasertib was a tolerable dosage.** Two-way ANOVA followed by Tukey test for multiple comparisons was used to compare the different treatment groups. Statistical comparison at the endpoint of control group was shown. VEHL: vehicle; CARBO: carboplatin; ns, not significant.

**Supplemental Figure S12. The mRNA level and correlation of PLK2 and PLK1 in carboplatin-treated patients of BrighTNess clinical trial. (A)** PLK2 mRNA expression is lower in tumors from TNBC patients who achieved a pathological complete response (pCR) as compared to tumors from patients with residual disease (RD). **(B)** PLK1 mRNA level is significantly higher in pCR than RD. **(C)** The negative correlation between PLK1 and PLK2 is more dramatic in pCR (r=-0.335) than RD (r=-0.13). TNBC patients treated with paclitaxel followed by doxorubicin and cyclophosphamide plus carboplatin with or without veliparib were analyzed. Veliparib addition didn't appear to offer a significant advantage and therefore included in the analysis to increase the statistical power.

**Supplemental Figure S13. Chromosome 5q genes PLK2, MAP3K1, RAD17, PIK3R1 CNV in breast cancer cell lines (A) and PDX models (B).** Note that overall breast cancer cell lines had a lower level (less blue) of 5q genes loss compared to PDX models, suggesting *in vitro* cultures of breast cancer cell lines may selectively attenuate the chromosome 5q loss.

**Supplemental Figure S14. Doxycycline-induced PLK2 expression reduced the therapeutic effect of volasertib in human BT20 xenografts. (A)** qPCR analysis showed that PLK2 mRNA expression was induced by doxycycline (DOX) water in the human BT20-iPLK2 tumors. n = 3 for BT20-iPLK2 control group. n = 4 for BT20-iPLK2 DOX group. Statistical significance was determined by unpaired student’s t-test. *, $p < 0.05$. **(B)** Western blot analysis and quantification showed that PLK2 protein expression was induced by doxycycline water (DOX) in the human BT20-iPLK2 tumors. n = 3 per group. Statistical significance was determined by unpaired student’s t-test. *, $p < 0.05$. **(C)** Tumor growth curves of human BT20-iPLK2 tumors under control and volasertib treatment with or without DOX. Volasertib treatment significantly reduced tumor growth for non-DOX treated tumors but not DOX treated tumors at the experimental endpoint. Tumor size was measured using a digital caliper twice a week. Two-way ANOVA followed by Tukey test for multiple comparisons was used to analyze the growth curves. Statistical comparison of the experimental endpoint was shown. Green arrow indicates when the doxycycline treatment began. Black arrows indicate when the mice were treated with drugs. DOX: doxycycline; VEHL: vehicle; ns, not significant; ****, $p < 0.0001$. **(D)** Two-way ANOVA analysis of log2 fold change (FC) of tumor volume starting from drug treatment to the end of experiment showed a reduced but not significant effect of volasertib in DOX-treated BT20-iPLK2 tumors due to the low mice number per group and high variation of the tumor growth. **(E)** Body weight of BT20-iPLK2 model suggested 25 mg/kg/wk of volasertib was a tolerable dosage and doxycycline didn’t have an impact on NSG mouse body weight. Two-way ANOVA followed by Tukey test for multiple comparisons was used to compare the different treatment groups. Statistical comparison of the experimental endpoint was shown. Green arrow indicates when the doxycycline treatment began. Black arrows indicate when the mice were treated with drugs. VEHL: vehicle; ns, not significant.

**Supplemental Figure S15. Doxycycline-induced PLK2 expression abolished the therapeutic effect of carboplatin and volasertib in mouse Plk2$^{-/-}$; p53$^{-/-}$ 1963B basal-like breast tumors. (A)** 1963B-iPLK2 model without DOX administration. **(B)** 1963B-iPLK2 model with DOX induction. Note that the therapeutic effect of carboplatin, volasertib, and combination treatment was abolished by DOX-induced PLK2 expression. **(C and D)** body weight measurement indicated 25 mg/kg/wk of volasertib was a tolerable dosage. Tumor size was measured using a caliper twice a week until control groups reached ~1500 mm$^3$. Two-way ANOVA followed by Tukey test for multiple comparisons was used. Statistical comparison of the experimental endpoint was shown. Green arrows indicate when the DOX treatment began. Black arrows indicate when the mice were treated with drugs. VEHL: vehicle; CARBO: carboplatin; ns, not significant; ****, $p < 0.0001$.

**Supplemental Figure S1.**

**A** PLK2 (chr5:57749809-57755966)

| Cancer Subset | In Peak? | Nearest Peak (click link to launch IGV) | #Genes in Peak | Q-value | Frequency of Deletion | | |
|---|---|---|---|---|---|---|---|
| | | | | | Overall | Focal | High-level |
| all_cancers | No | chr5:58261529-59787458 | 2 | 4.96E-52 | 0.2515 | 0.0608 | 0.0328 |
| Breast cancer | Yes | chr5:54834826-57786659 | 11 | 1.37E-4 | 0.2294 | 0.0493 | 0.0287 |
| Lung adenocarcinoma | Yes | chr5:914233-180360469 | 1019 | 0.0048 | 0.2773 | 0.0532 | 0.0168 |
| Acute myeloid leukemia | Yes | chr5:42547255-58652895 | 59 | 1.0 | 0.0155 | 0.0052 | 0.0 |
| Ovarian serous carcinoma | No | chr5:58147456-59787458 | 2 | 9.55E-41 | 0.5506 | 0.2469 | 0.1439 |
| Uterine and endometrial carcinoma | No | chr5:58261529-59787458 | 2 | 1.89E-10 | 0.1774 | 0.0605 | 0.0222 |
| Head and neck squamous cell carcinoma | No | chr5:58261529-59787458 | 2 | 7.86E-4 | 0.371 | 0.0613 | 0.0226 |
| Colorectal cancer | No | chr5:58261529-59787458 | 2 | 0.0207 | 0.1829 | 0.0274 | 0.0103 |
| Lung squamous cell carcinoma | No | chr5:58261529-59787458 | 2 | 0.0635 | 0.686 | 0.0523 | 0.0523 |
| Bladder cancer | No | chr5:58261529-59787458 | 2 | 0.118 | 0.4265 | 0.0809 | 0.0294 |
| Glioblastoma multiforme | No | chr5:158528732-180360469 | 181 | 1.0 | 0.0293 | 0.0034 | 0.0052 |
| Kidney clear cell carcinoma | No | No peak on chromosome | 0 | 1.0 | 0.0161 | 0.0040 | 0.0020 |

| Legend: Cancer-Promoter Significance | |
|---|---|
| q <= 0.25 | In a peak |
| q <= 0.25 | Not in a peak |
| q > 0.25 | In a peak |
| q > 0.25 | Not in a peak |

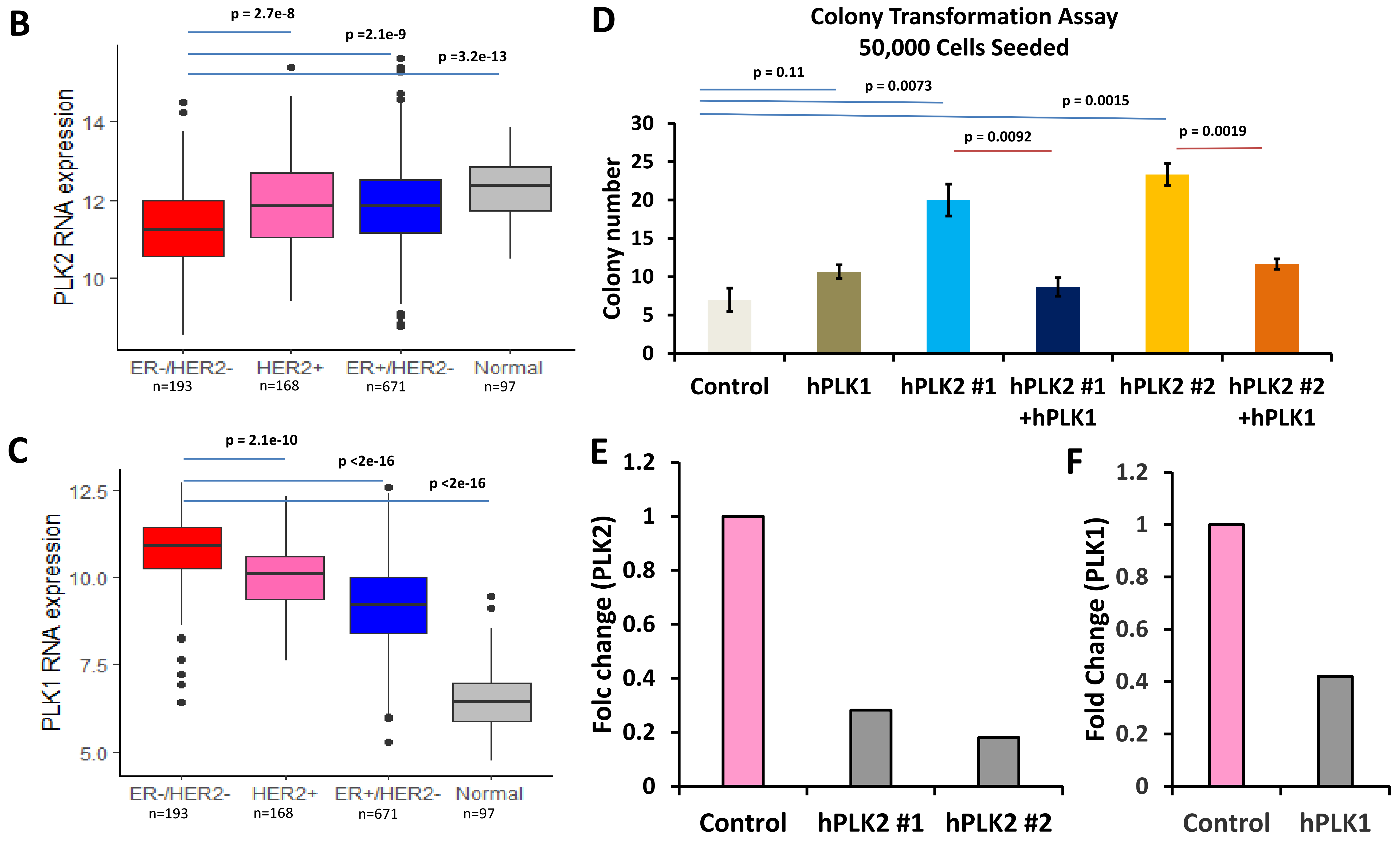

**Supplemental Figure S2.**

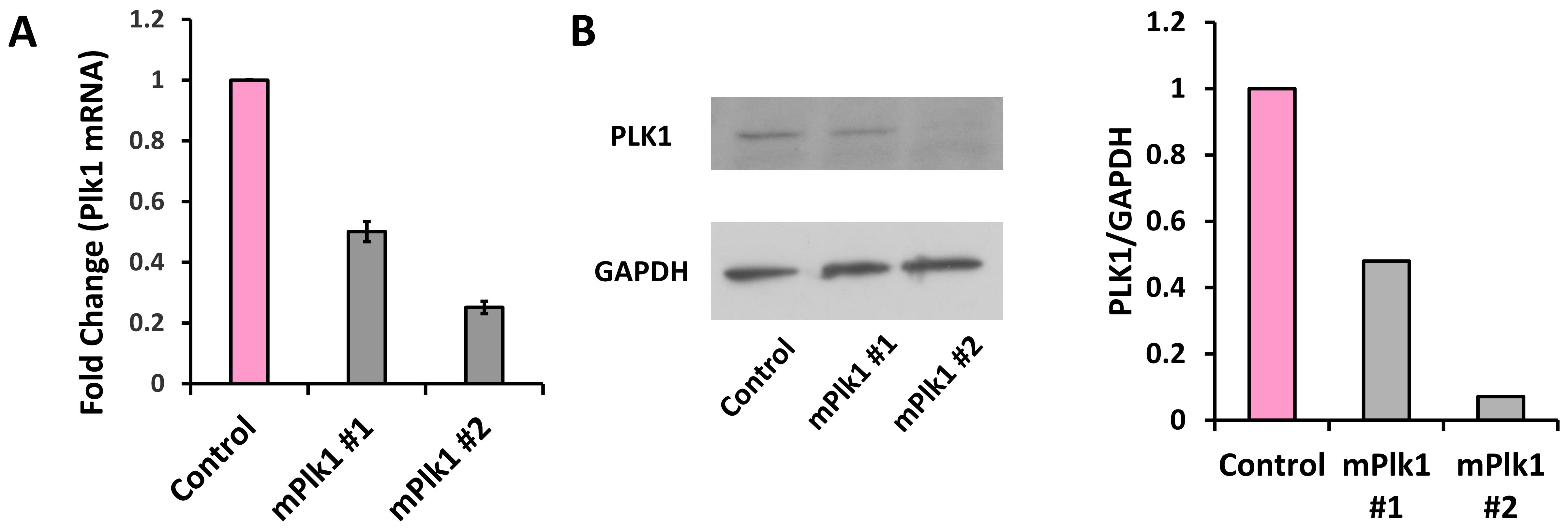

**Supplemental Figure S3.**

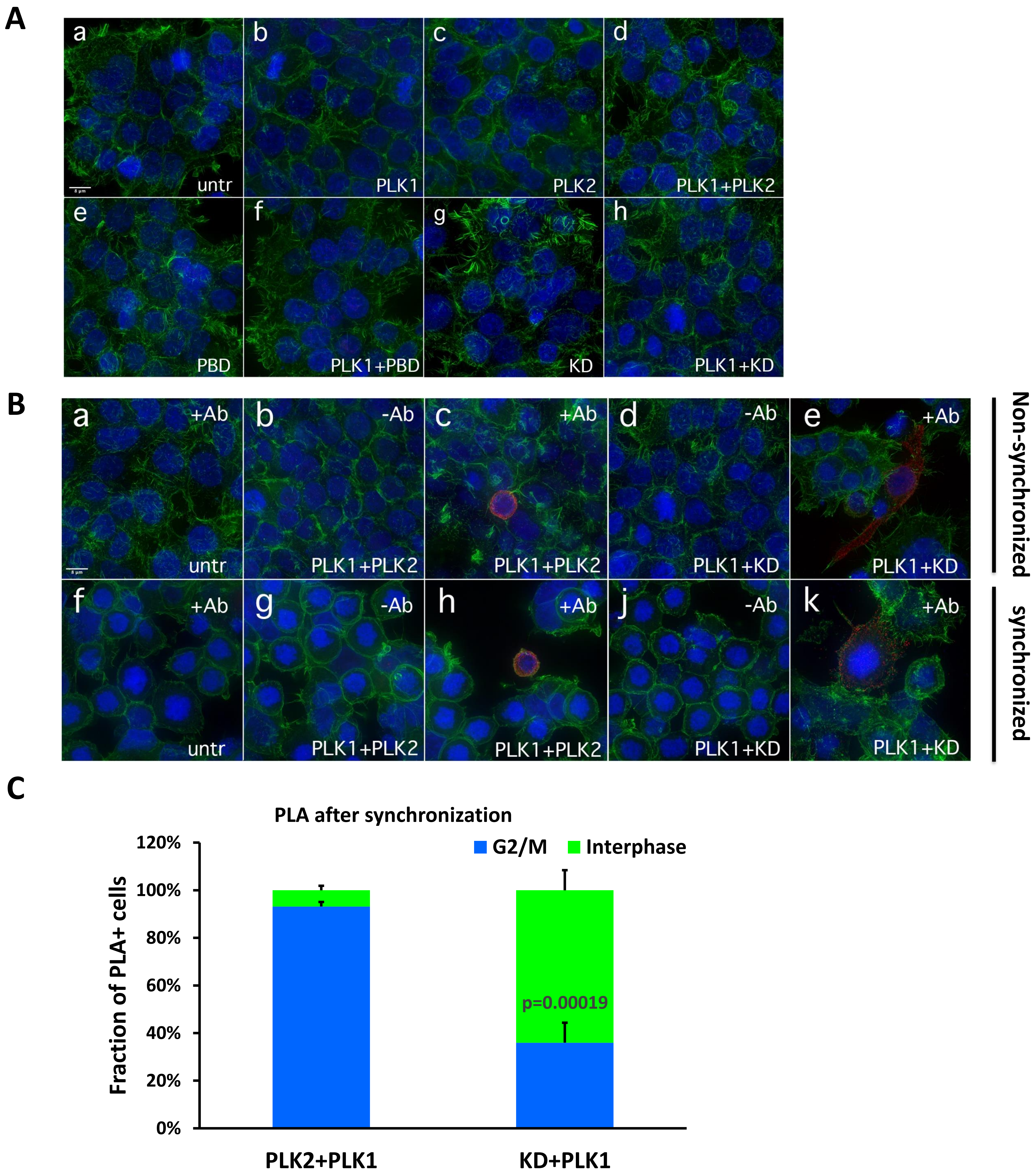

**Supplemental Figure S4.**

A

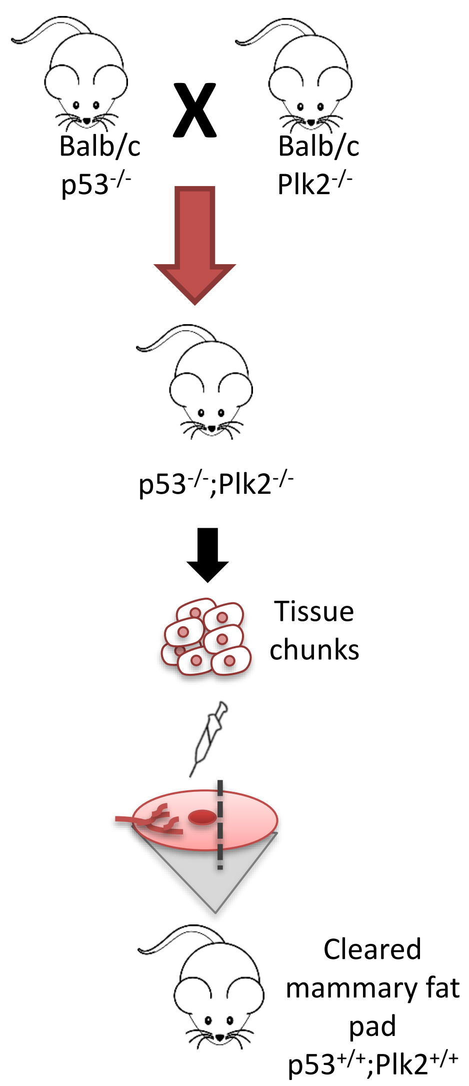


B

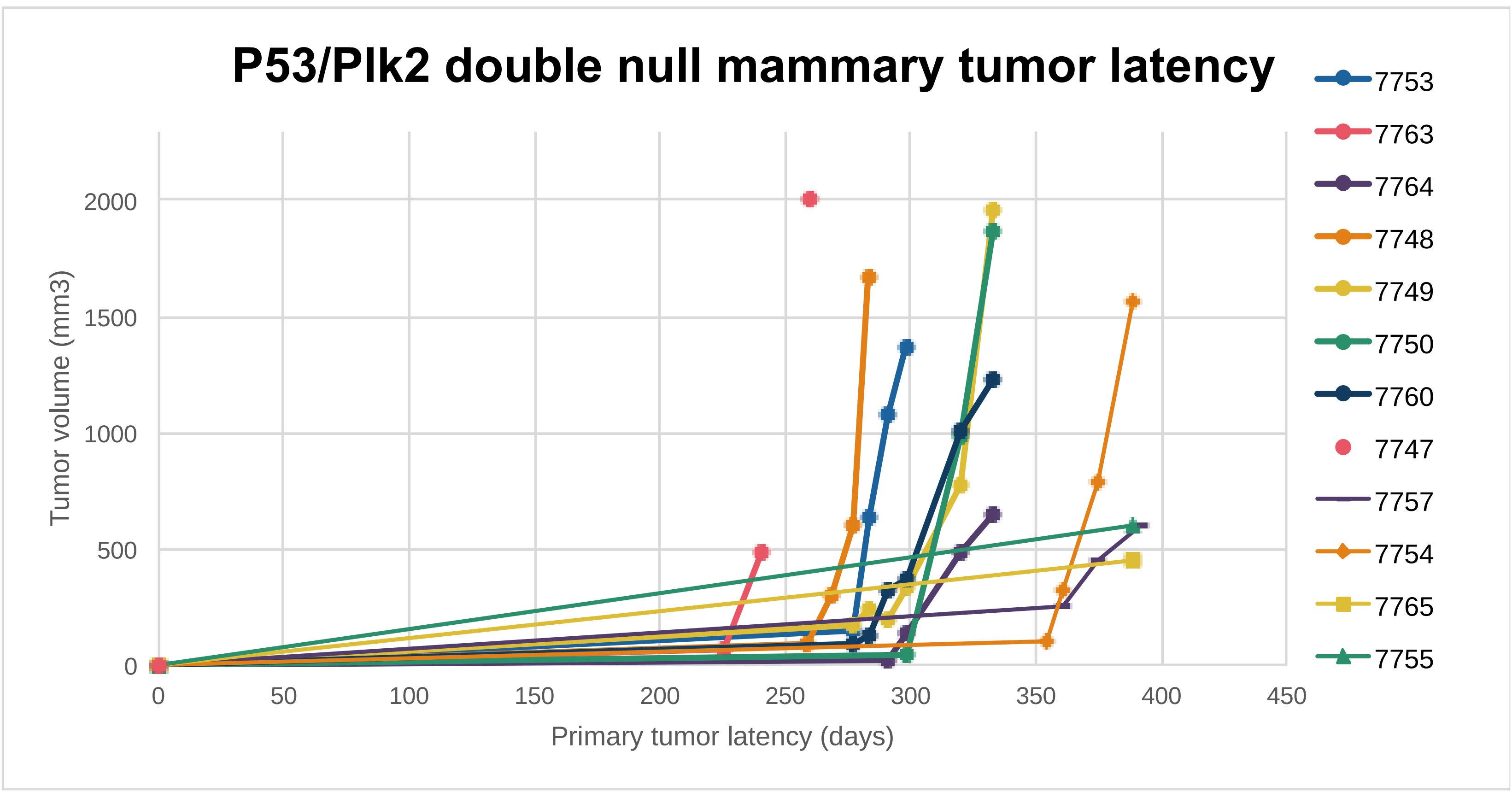

**Supplemental Figure S5.**

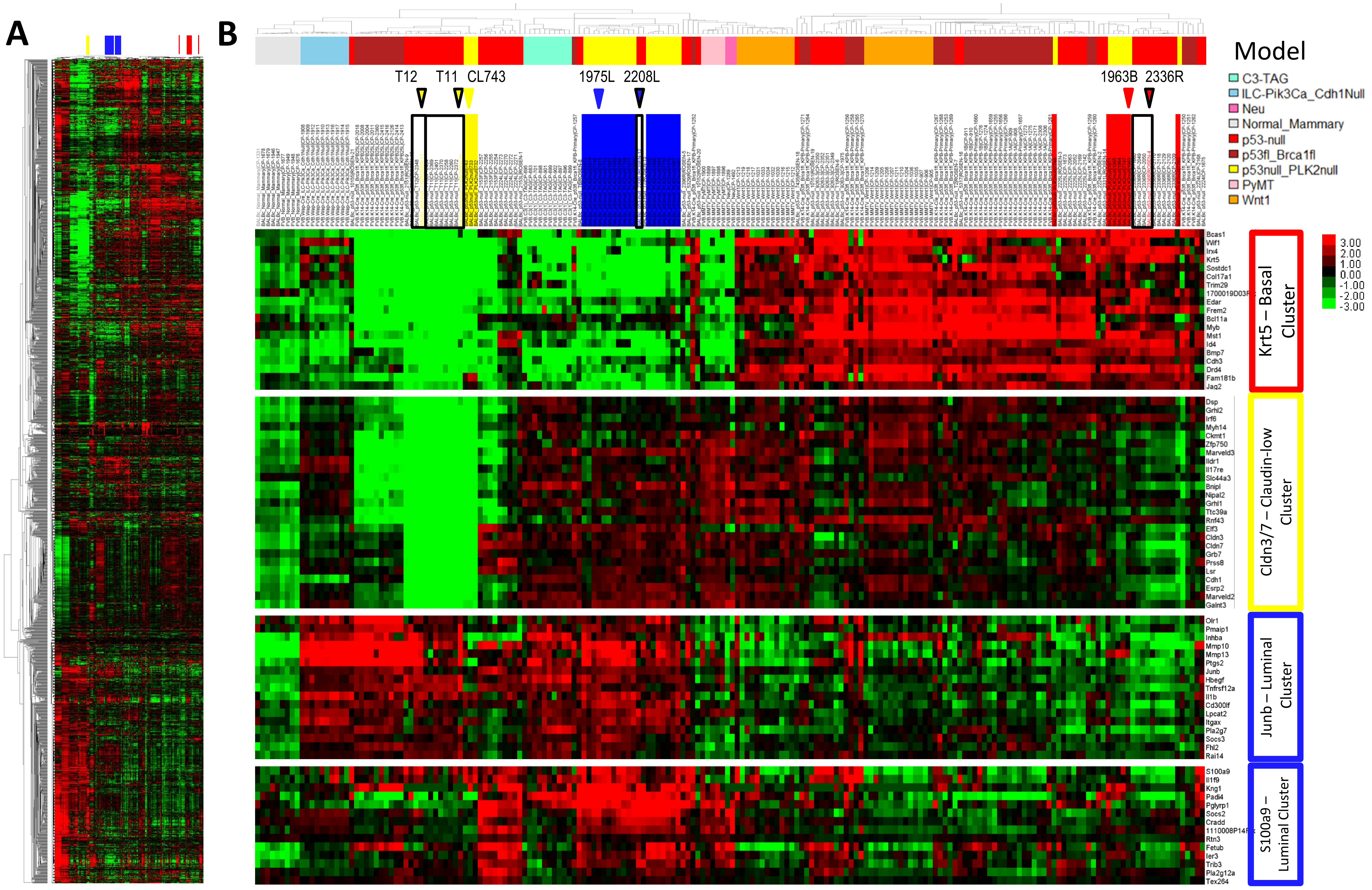

**Supplemental Figure S6.**

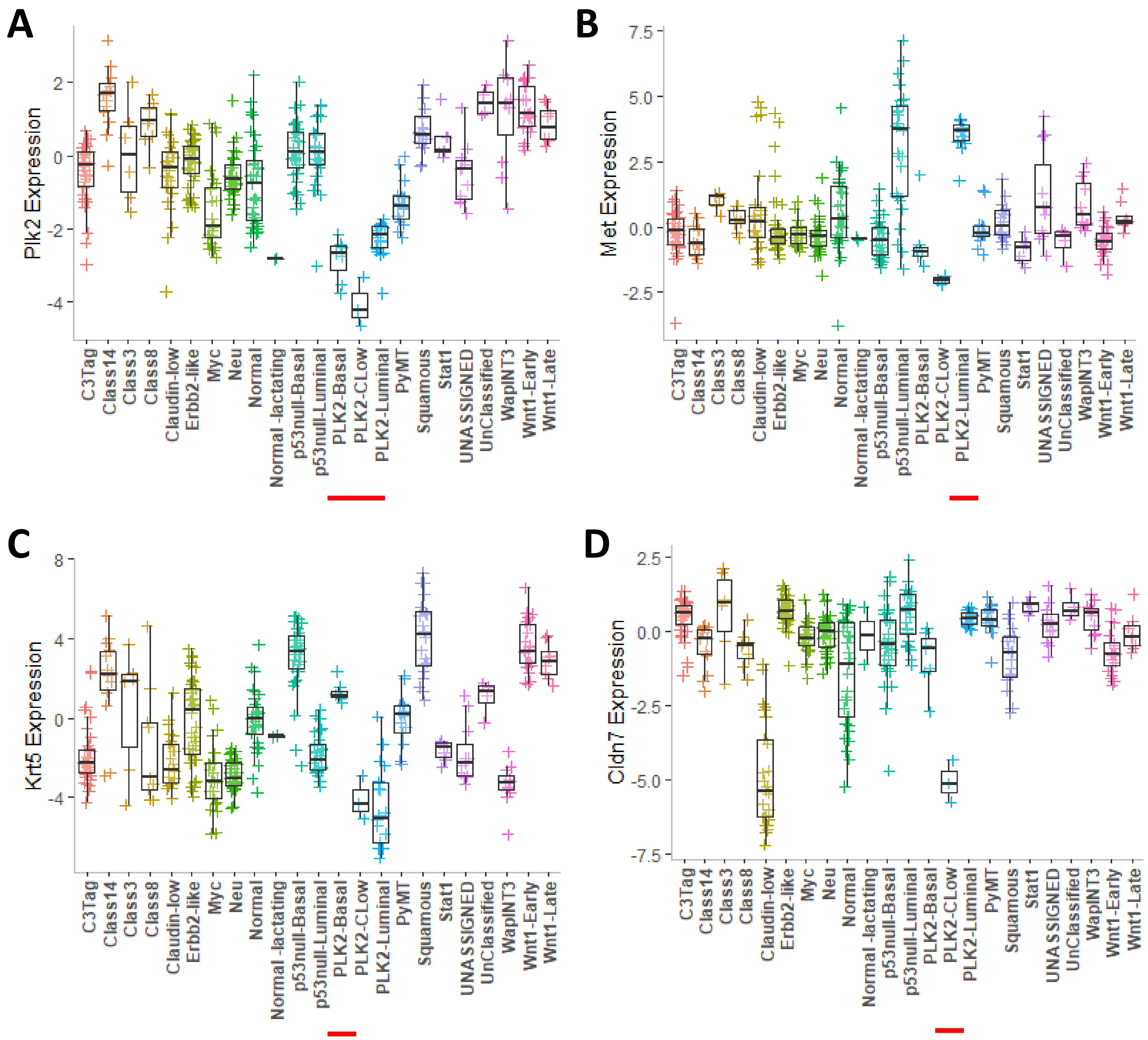

**Supplemental Figure S7.**

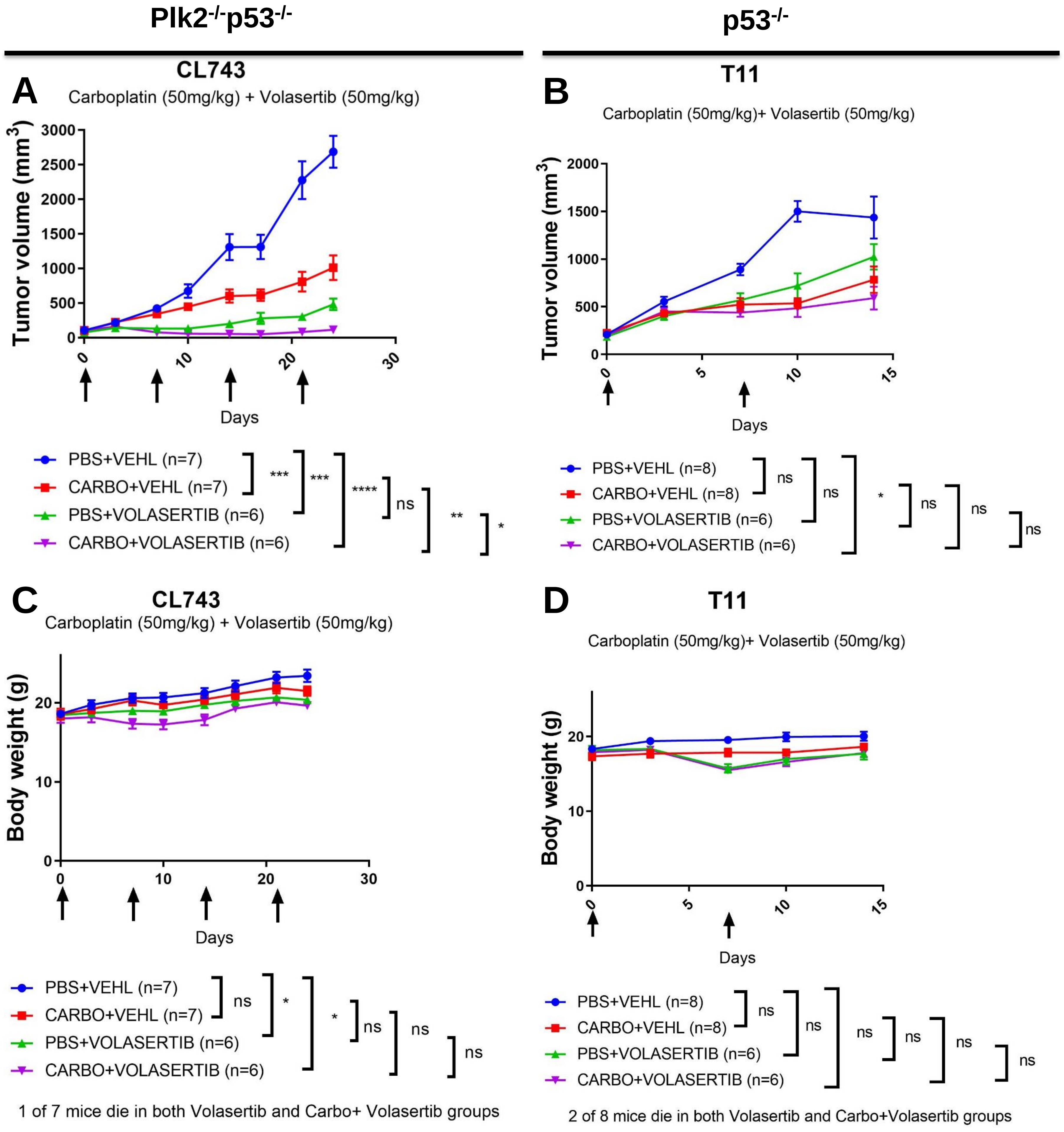

**Supplemental Figure S8.**

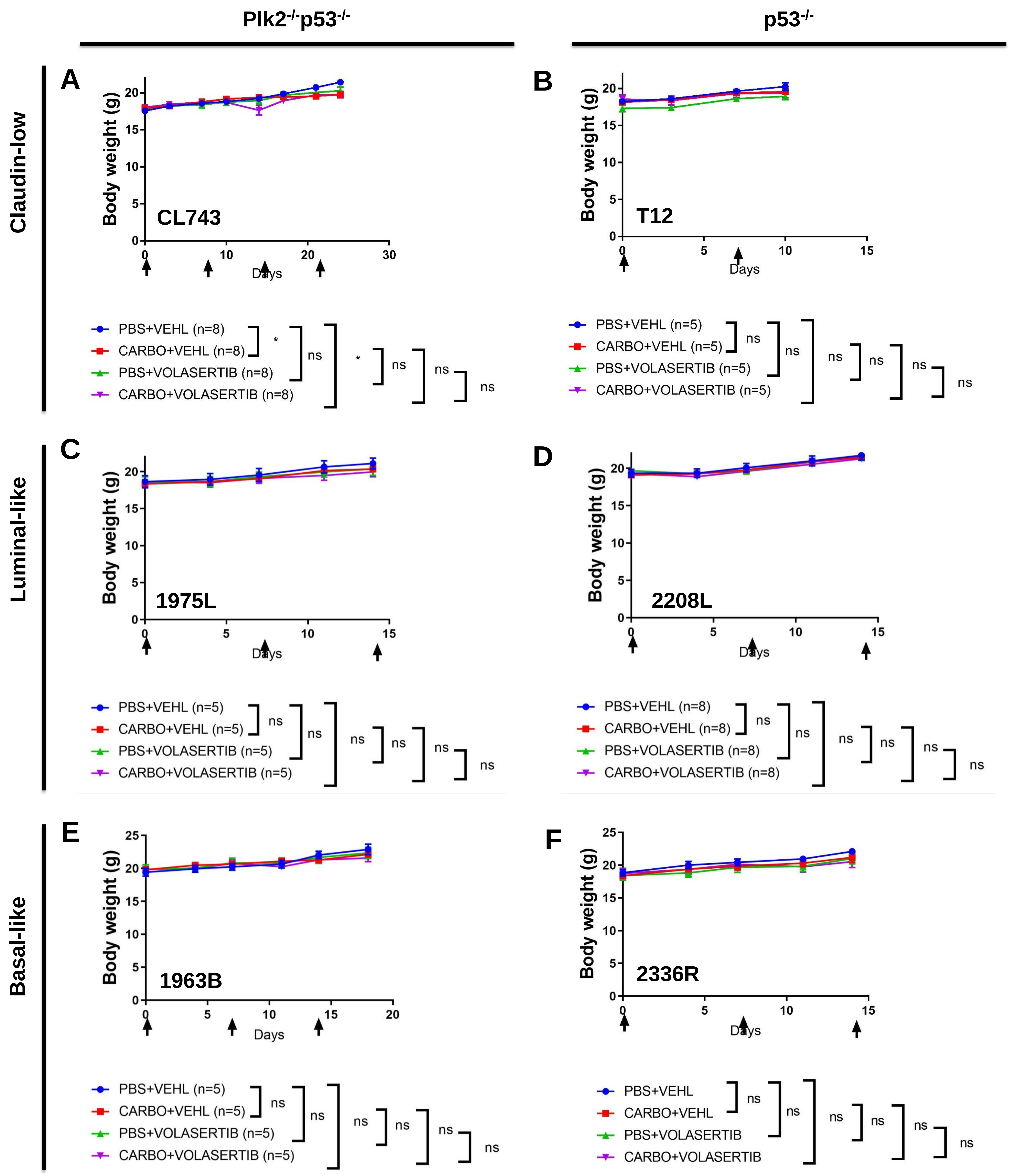

**Supplemental Figure S9.**

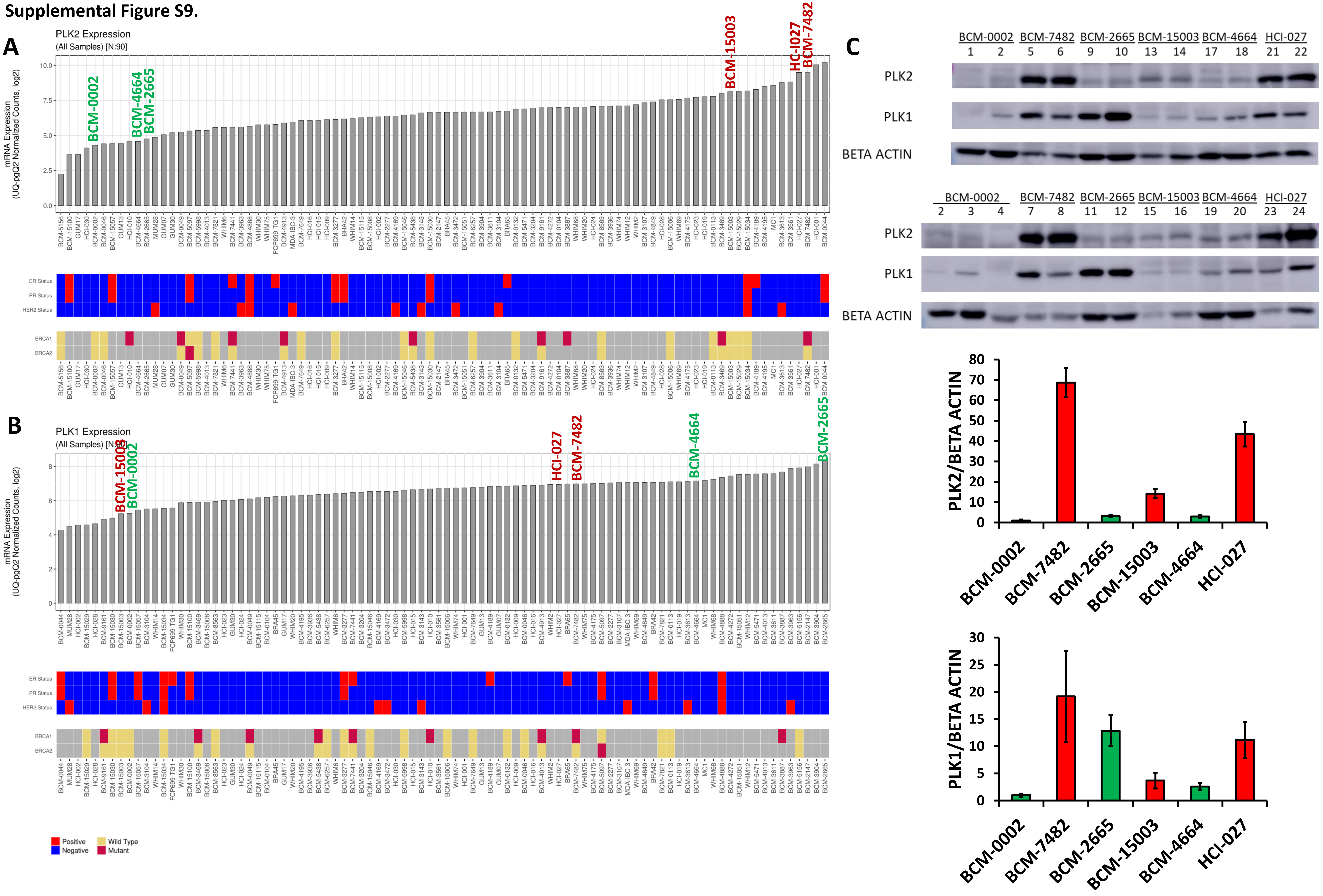

**Supplemental Figure S10.**

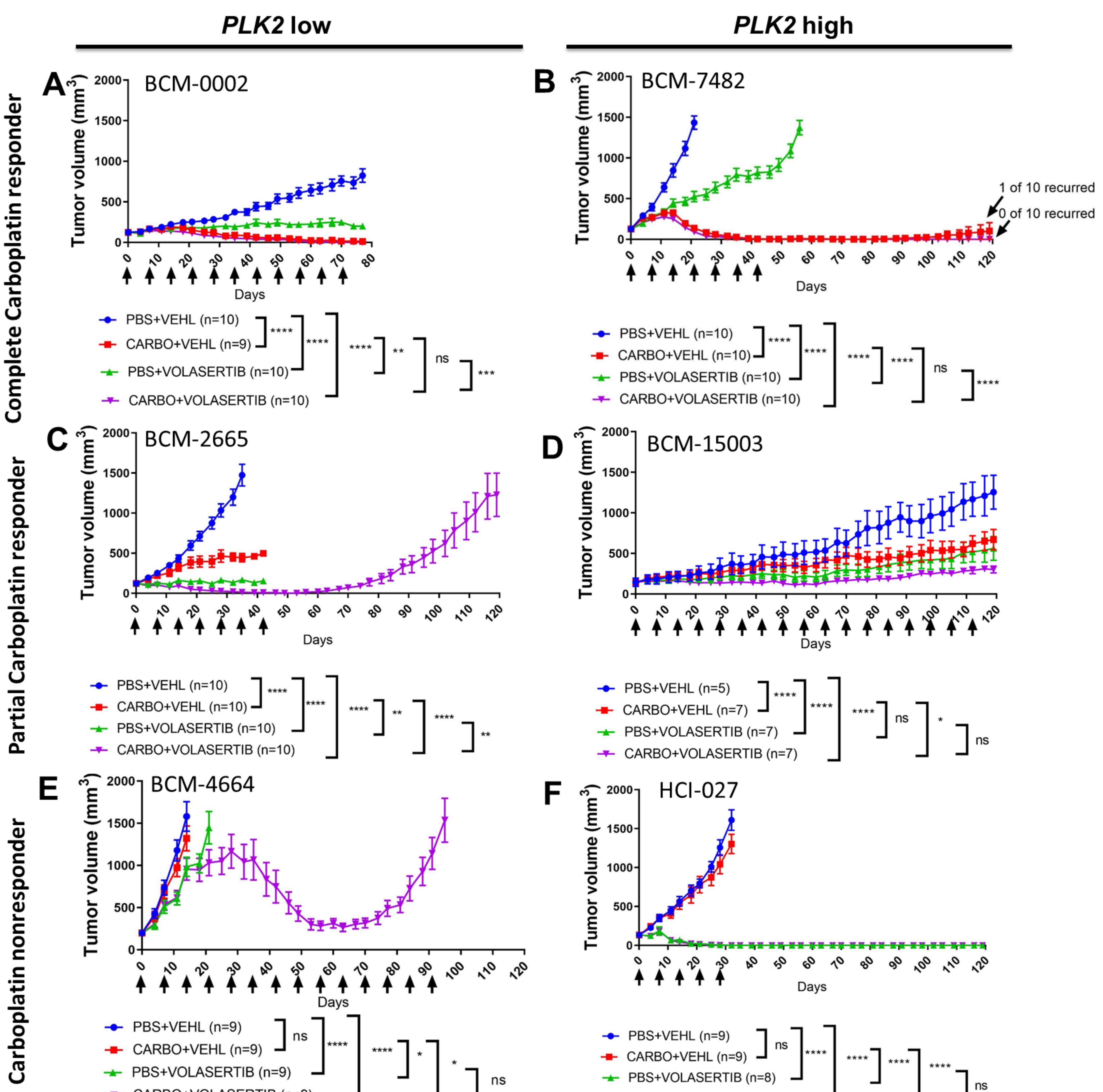

Supplemental Figure S11.

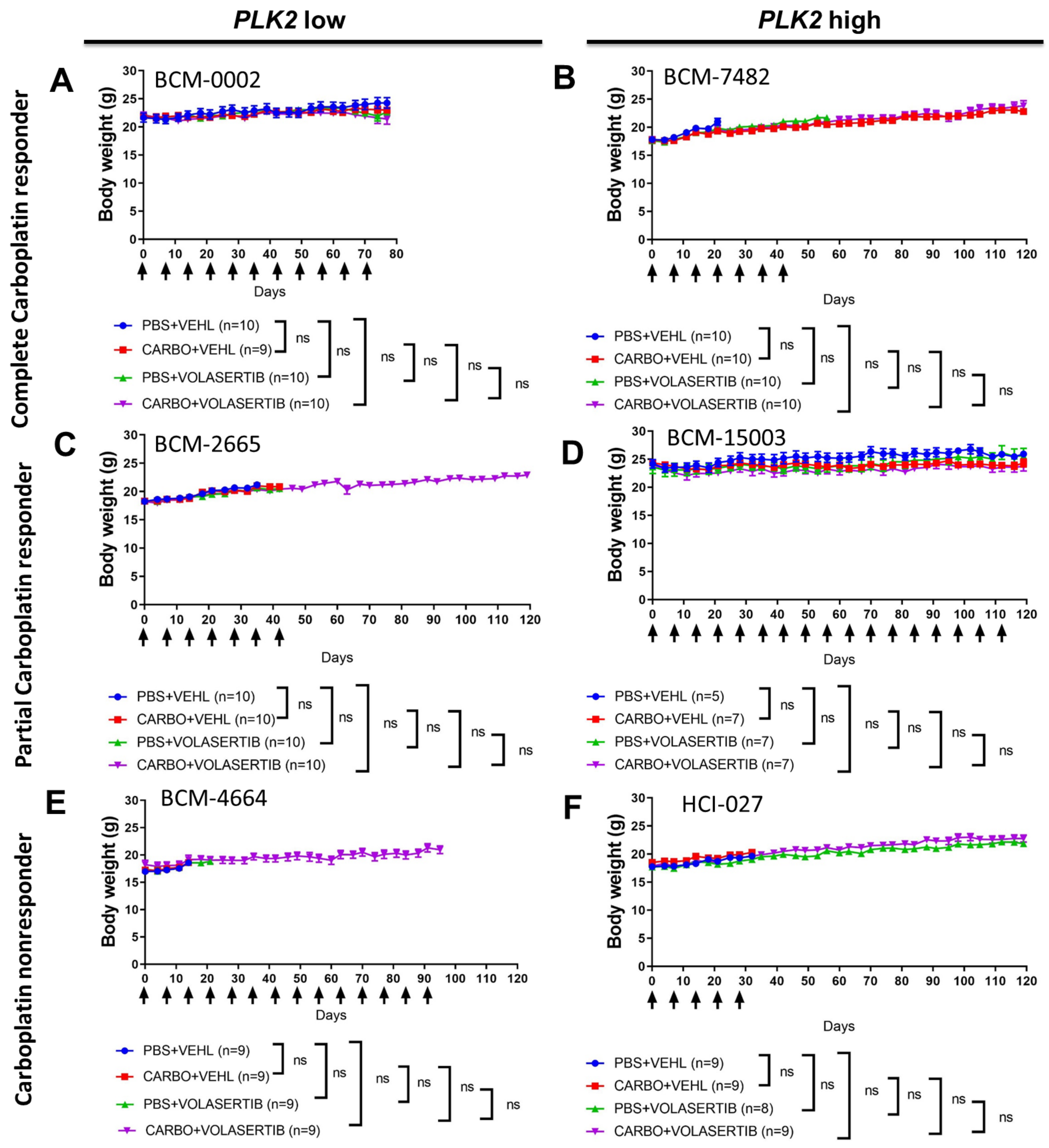

**Supplemental Figure S12.**

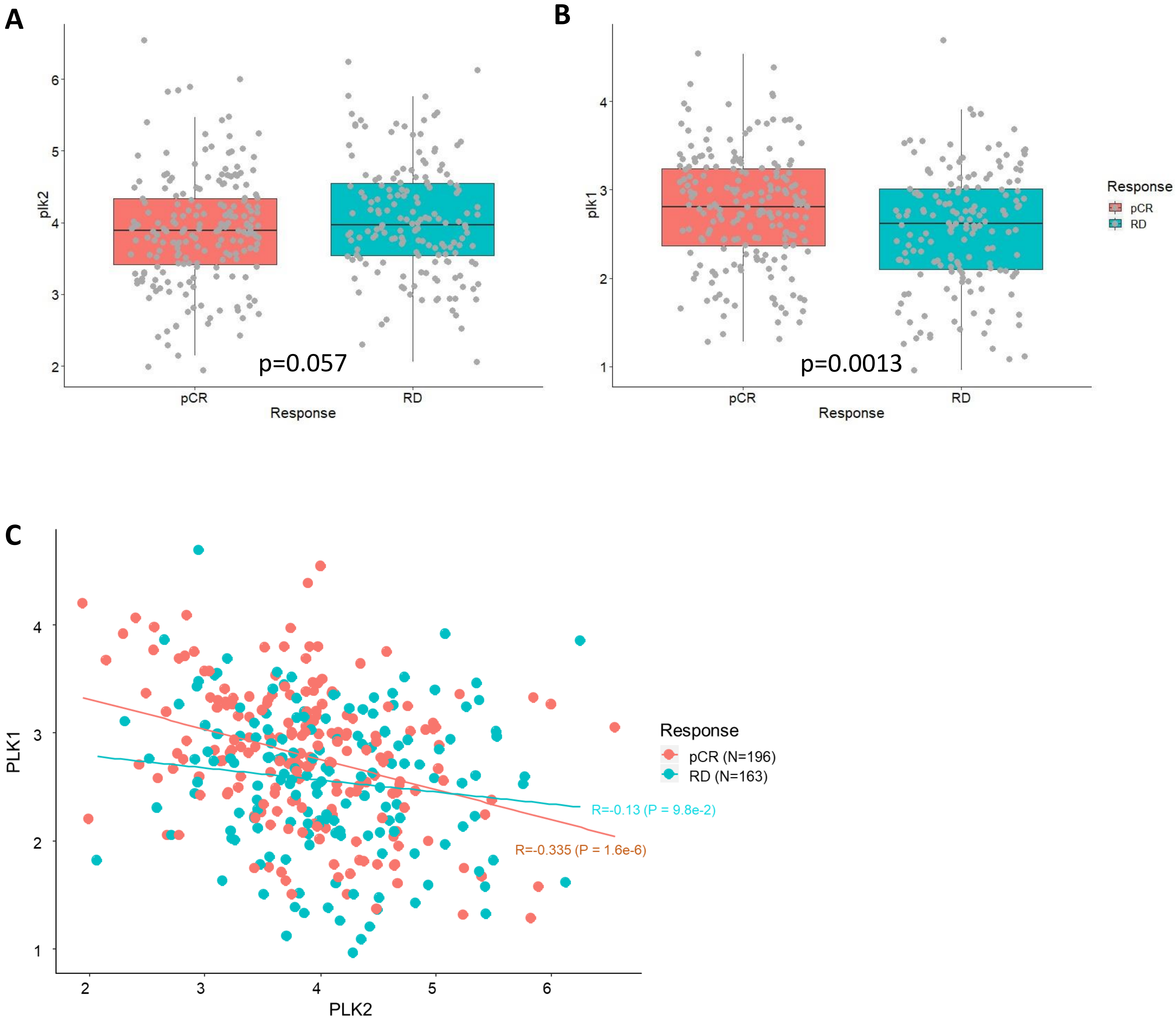

**Supplemental Figure S13.**

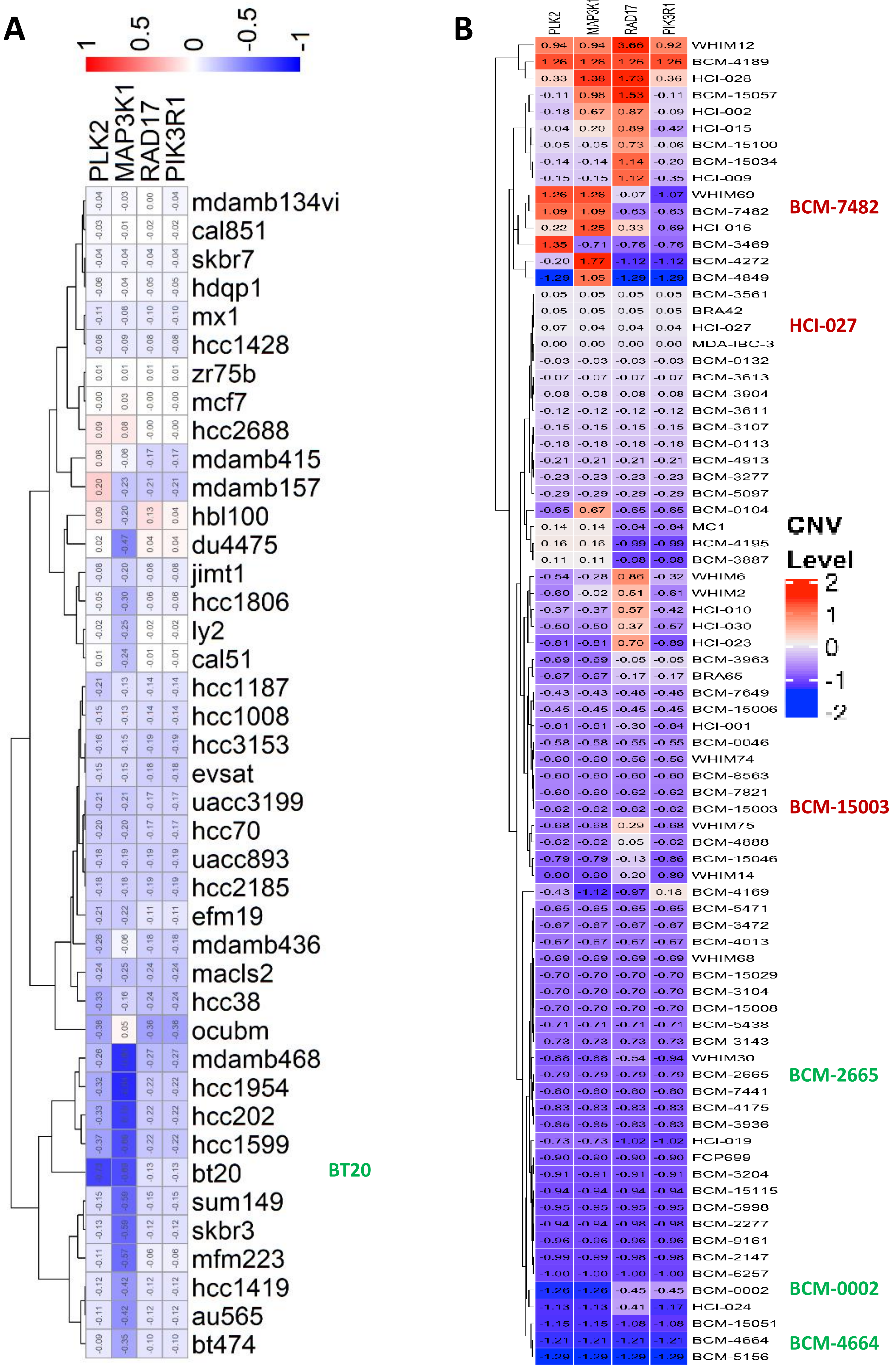

Supplemental Figure S14.

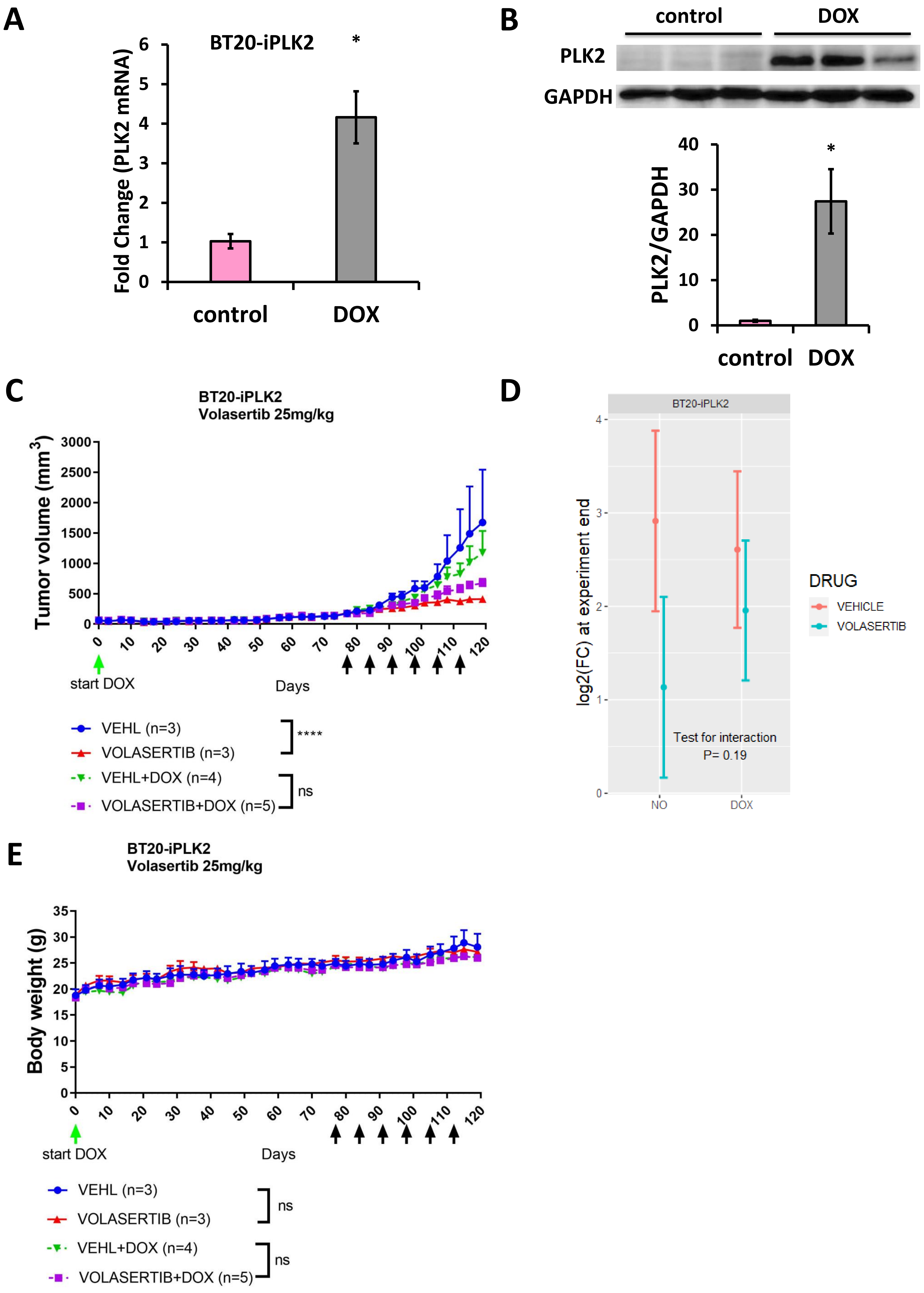

**Supplemental Figure S15.**

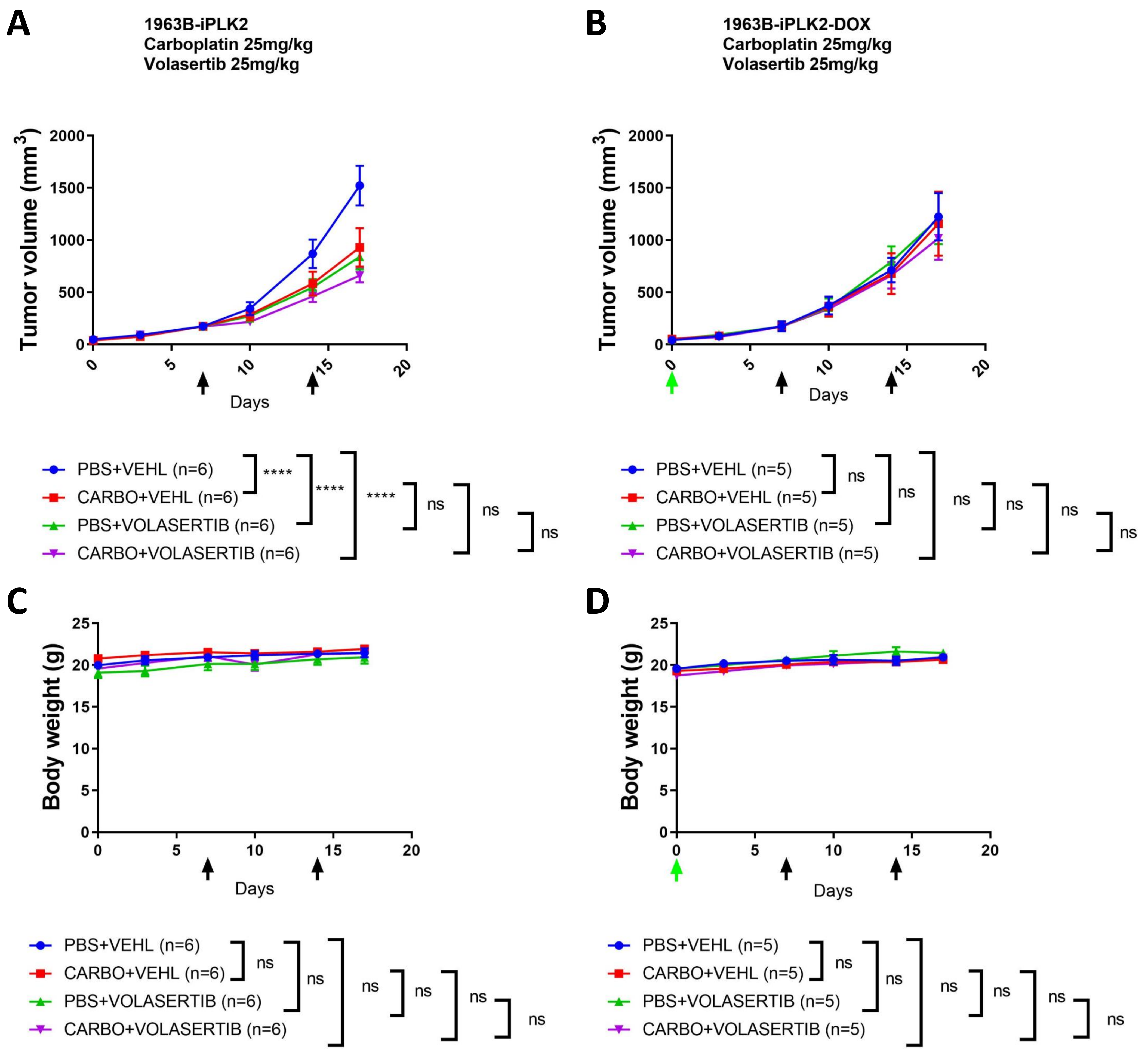

**Supplemental Table S1.**

| Genotype | Number | Percentage (%) | Theoretical percentage (%) |
|---|---|---|---|
| **$Plk2^{+/-}$; $Plk1^{+/-}$ male X $Plk2^{+/-}$; $Plk1^{+/-}$ female<br>Total number of pups: 91<br>Total number of litters: 20<br>Average number of pups per litter: 4.55** | | | |
| Genotype | Number | Percentage (%) | Theoretical percentage (%) |
| $Plk2^{-/-}$; $Plk1^{+/+}$ | 4 | 4.40 | 6.25 |
| $Plk2^{+/-}$; $Plk1^{+/+}$ | 20 | 21.98 | 12.50 |
| $Plk2^{+/+}$; $Plk1^{+/+}$ | 22 | 24.18 | 6.25 |
| $Plk2^{-/-}$; $Plk1^{+/-}$ | 0 | 0 | 12.50 |
| $Plk2^{+/-}$; $Plk1^{+/-}$ | 25 | 27.47 | 25.00 |
| $Plk2^{+/+}$; $Plk1^{+/-}$ | 20 | 21.98 | 12.50 |
| $Plk2^{-/-}$; $Plk1^{-/-}$ * | 0 | 0 | 6.25 |
| $Plk2^{+/-}$; $Plk1^{-/-}$ * | 0 | 0 | 12.50 |
| $Plk2^{+/+}$; $Plk1^{-/-}$ * | 0 | 0 | 6.25 |
| **$Plk2^{+/-}$; $Plk1^{+/-}$ male X $Plk2^{-/-}$ female<br>Total number of pups: 50<br>Total number of litters: 15<br>Average number of pups per litter: 3.33** | | | |
| Genotype | Number | Percentage (%) | Theoretical percentage (%) |
| $Plk2^{-/-}$; $Plk1^{+/+}$ | 3 | 6.00 | 25.00 |
| $Plk2^{+/-}$; $Plk1^{+/+}$ | 23 | 46.00 | 25.00 |
| $Plk2^{-/-}$; $Plk1^{+/-}$ | 0 | 0 | 25.00 |
| $Plk2^{+/-}$; $Plk1^{+/-}$ | 24 | 48.00 | 25.00 |
| **$Plk2^{+/-}$; $Plk1^{+/-}$ male X $Plk2^{+/-}$ female<br>Total number of pups: 34<br>Total number of litters: 7<br>Average number of pups per litter: 4.86** | | | |
| Genotype | Number | Percentage (%) | Theoretical percentage (%) |
| $Plk2^{-/-}$; $Plk1^{+/+}$ | 1 | 2.94 | 12.50 |
| $Plk2^{+/-}$; $Plk1^{+/+}$ | 8 | 23.53 | 25.00 |
| $Plk2^{+/+}$; $Plk1^{+/+}$ | 8 | 23.53 | 12.50 |
| $Plk2^{-/-}$; $Plk1^{+/-}$ | 0 | 0 | 12.50 |
| $Plk2^{+/-}$; $Plk1^{+/-}$ | 11 | 32.35 | 25.00 |
| $Plk2^{+/+}$; $Plk1^{+/-}$ | 6 | 17.65 | 12.50 |

* Note that $Plk1^{-/-}$ is embryonic lethal but $Plk1^{+/-}$ and $Plk2^{-/-}$ are viable

**Supplemental Table S2.**

| Luminal-like TNBC | | Basal-like TNBC | | Claudin-low TNBC | |
|---|---|---|---|---|---|
| $Plk2^{-/-}$; $p53^{-/-}$ | $p53^{-/-}$ | $Plk2^{-/-}$; $p53^{-/-}$ | $p53^{-/-}$ | $Plk2^{-/-}$; $p53^{-/-}$ | $p53^{-/-}$ |
| 1975L | 2208L | 1963B | 2336R | CL743 | T11, T12 |

**Supplemental Table S3.**

| Species | Name | Cat# | Clone ID | Mature Antisense |
|---|---|---|---|---|
| mouse | mPlk1 #1 | RMM3981-201757953 | TRCN0000027646 | ATAAGTTTGGTGTGGTCCTGG |
| mouse | mPlk1 #2 | RMM3981-201757918 | TRCN0000027611 | TTTATTGAGGACTTTGAGAGG |
| human | hPLK1 | RHS4430-200284623 | V3LHS_311461 | CTGTCTGAAGCATCTTCTG |
| human | hPLK2 #1 | RHS4430-99880311 | V3LHS_634296 | TTAATTCTGATGAACAGCC |
| human | hPLK2 #2 | RHS4430-98893790 | V2LHS_91309 | TAAAGAGCATCATTAGGGC |
